# Cdc25-Mediated Activation of the Small GTPase RasB Is Essential for Hyphal Fusion and Symbiotic Infection of *Epichloë festucae*

**DOI:** 10.1101/2025.11.03.686139

**Authors:** Mariko Inagaki, Shota Kamiya, Ayane Okamura, Atsushi Miura, Yuka Kayano, Aiko Tanaka, Daigo Takemoto

**Author notes:** **Corresponding Author**: Daigo Takemoto. These authors contributed equally to this work (Mariko Inagaki, Shota Kamiya).

## Abstract

*Epichloë festucae* is a filamentous endophytic fungus that symbiotically colonizes the intercellular spaces of aerial tissues in perennial ryegrass (*Lolium perenne*) without causing disease symptoms. This mutualistic association enhances host resistance to both biotic and abiotic stresses. Balanced and coordinated growth of *E. festucae* with its host is essential for the establishment and long-term maintenance of the symbiotic relationship. Various *E. festucae* mutants defective in symbiosis with host plants have been isolated, and notably, many of these symbiosis-defective mutants also lack hyphal fusion ability under culture conditions, supporting a close functional connection between signal transduction required for hyphal fusion and symbiosis establishment. Using a plasmid insertional mutagenesis approach, we identified Cdc25 as an essential regulator of hyphal fusion in *E. festucae*. Cdc25 encodes a guanine nucleotide exchange factor (GEF) that activates the small GTPase Ras. The *Δcdc25* strain lost both hyphal fusion ability and the capacity to infect host plants. Furthermore, yeast two-hybrid assays revealed that Cdc25 specifically interacts with RasB, one of the five Ras proteins in *E. festucae*. Expression of constitutively active (CA) RasB in the *Δcdc25* strain restored both hyphal fusion and host infection, whereas expression of CA-RasB in the *ΔmpkB* deletion strain failed to rescue its defect in hyphal fusion, suggesting that the Cdc25-RasB signaling module acts upstream of the MAPK cascade. In pathogenic fungi, this signaling module is known to regulate infection-related morphogenesis. These findings indicate that *E. festucae* has evolutionarily repurposed the conserved Cdc25-RasB module to coordinate hyphal fusion and maintain a stable mutualistic interaction with its host.

## 1 Introduction

*Epichloë festucae* is a filamentous fungal endophyte that forms a symptomless mutualistic association with cool-season grasses such as perennial ryegrass (*Lolium perenne*). Within aerial host tissues, the fungus grows intercellularly without inducing disease symptoms, maintaining a regulated balance between fungal growth and host development (Christensen et al., 2008; Tanaka et al., 2012). Colonization by endophytic fungi provides multiple benefits to the host plant, enhancing its fitness under various environmental stresses (Schardl et al., 2004; Kuldau and Bacon, 2008). In host plants, *Epichloë* endophytes synthesize several classes of biologically active metabolites that strengthen plant resistance to diverse stresses, including drought, microbial diseases, and insect or mammalian herbivory (Rowan et al., 1986; Bush et al., 1997; Wilkinson et al., 2000; Tanaka et al., 2005; Saikia et al., 2012; Niones and Takemoto, 2015).

In *E. festucae*, a number of genes essential for establishing symbiotic infection have been identified. The first gene shown to be required for mutualistic colonization was *noxA*, which encodes an NADPH oxidase responsible for reactive oxygen species (ROS) production (Tanaka et al., 2006). The *noxA* mutant exhibits unrestricted hyphal growth within host tissues, resulting in severe stunting and often premature death of the host plant. Subsequent studies demonstrated that NoxA activity depends on its regulatory components, NoxR (a human p67^phox^-like protein) and the small GTPase RacA, both of which are indispensable for maintaining a balanced endophytic symbiosis (Takemoto et al., 2006; Tanaka et al., 2008; Kayano et al., 2018). Interestingly, *noxA*, *noxR*, and *racA* mutants are also defective in hyphal cell-cell fusion, suggesting that ROS-mediated signaling is closely linked to the development of hyphal networks within host tissues (Kayano et al., 2013).

In addition to the Nox pathway, several other genes are implicated in both hyphal fusion and symbiotic infection. These include *proA*, which encodes a C6 zinc-finger transcription factor homologous to ADV-1 of *Neurospora crassa*, as well as *symB* (HAM-7) and *symC* (IDC-3) (Tanaka et al., 2013; Green et al., 2017). Mutants lacking these genes show severe growth defects in host plants, leading to stunting phenotypes similar to those observed in *noxA* and *noxR* mutants, implying that proper communication between hyphae is essential for controlled fungal proliferation during symbiosis.

Genes associated with the cell wall integrity (CWI) MAP kinase pathway have also been implicated in regulating symbiotic development. The MAP kinase MpkA (homologous to MAK-1 in *N. crassa*) and its scaffold protein SO are both required for hyphal fusion and successful symbiotic infection (Charlton et al., 2012; Becker et al., 2015). Another MAPK cascade centered on MpkB governs transcriptional programs essential for both hyphal fusion and *in planta* colonization. The nuclear protein NsiA interacts with MpkB and controls expression of genes required for symbiotic infection and hyphal fusion, thereby integrating MAPK signaling with intercellular communication (Tanaka et al., 2020). Loss of these components results in highly similar phenotypes-loss of hyphal fusion, impaired colonization, and collapse of mutualism, indicating that fungal cell-cell signaling is directly linked to the establishment of symbiotic growth.

Interestingly, most genes identified as essential for symbiotic development in *E. festucae* correspond to genes that are also required for hyphal fusion in the model fungus *N. crassa*, including MAPK components (MAK-2, MEK-2), small GTPases (RAC-1), NADPH oxidase subunits (NOX-1, NOR-1), the CWI scaffold SO, and the transcription factor PP-1 (Fischer and Glass, 2019). In *N. crassa*, these genes constitute a conserved cell-to-cell communication pathway that mediates chemotropic growth and fusion during colony formation. The conservation of these signaling modules suggests that *E. festucae* has evolutionarily co-opted ancestral fusion machinery to regulate its coordinated growth and communication within plant tissues.

To further understand the genetic basis of hyphal fusion and its connection to symbiotic development, we conducted a mutagenesis screen for *E. festucae* strains defective in hyphal fusion. Through this approach, we identified a candidate gene, Cdc25, which encodes a Ras-specific guanine nucleotide exchange factor (GEF). Given the established importance of small GTPases and MAPK signaling in regulating fungal morphogenesis and communication, Cdc25 was hypothesized to function as a key upstream regulator within the signaling network required for both hyphal fusion and mutualistic interaction in the endophyte. In this study, we investigated the role of Cdc25 and its downstream components to clarify how Ras-mediated signaling contributes to symbiotic infection in *E. festucae*.

## 2 Results

### 2.1 Isolation of Hyphal Fusion-Deficient Mutants of *E. festucae*

To investigate the signaling mechanisms that link hyphal fusion with the establishment of symbiosis, we aimed to isolate *E. festucae* mutant strains that had lost the ability to undergo hyphal fusion. Random insertional mutagenesis was performed using the restriction enzyme–mediated integration (REMI) method (Sánchez et al., 1998) with *Pst*I, in which the transformation vector pNPP1 (containing hygromycin resistance cassette and *GFP* genes) was introduced into protoplasts of *E. festucae* strain Fl1. Following transformation, strains showing GFP fluorescence were transferred to a nutrient-poor 3% agar medium and cultured for two weeks. Hyphae grown on 3% agar were stained with Calcofluor white and examined by fluorescence microscopy to identify strains lacking hyphal fusion. Among approximately 1,200 REMI transformants examined, three strains (RPA41, RPA112, and RPA519) exhibiting reduced or no hyphal fusion were isolated.

To compare the frequency of hyphal fusion among the wild-type strain, three REMI mutant strains, and the *Δso*, *ΔmpkB*, and *Δham8* knockout strains, which are known to be defective in hyphal fusion (Charlton et al., 2012; Tanaka et al., 2020), each strain was cultured on 3% agar medium. While frequent hyphal fusions were observed in the wild-type strain, no hyphal fusion was detected in strains RPA41, RPA112, *Δso*, *ΔmpkB* and *Δham8*. Only occasional hyphal fusion events were observed in strain RPA519 (Figure 1a).

**FIGURE 1.**
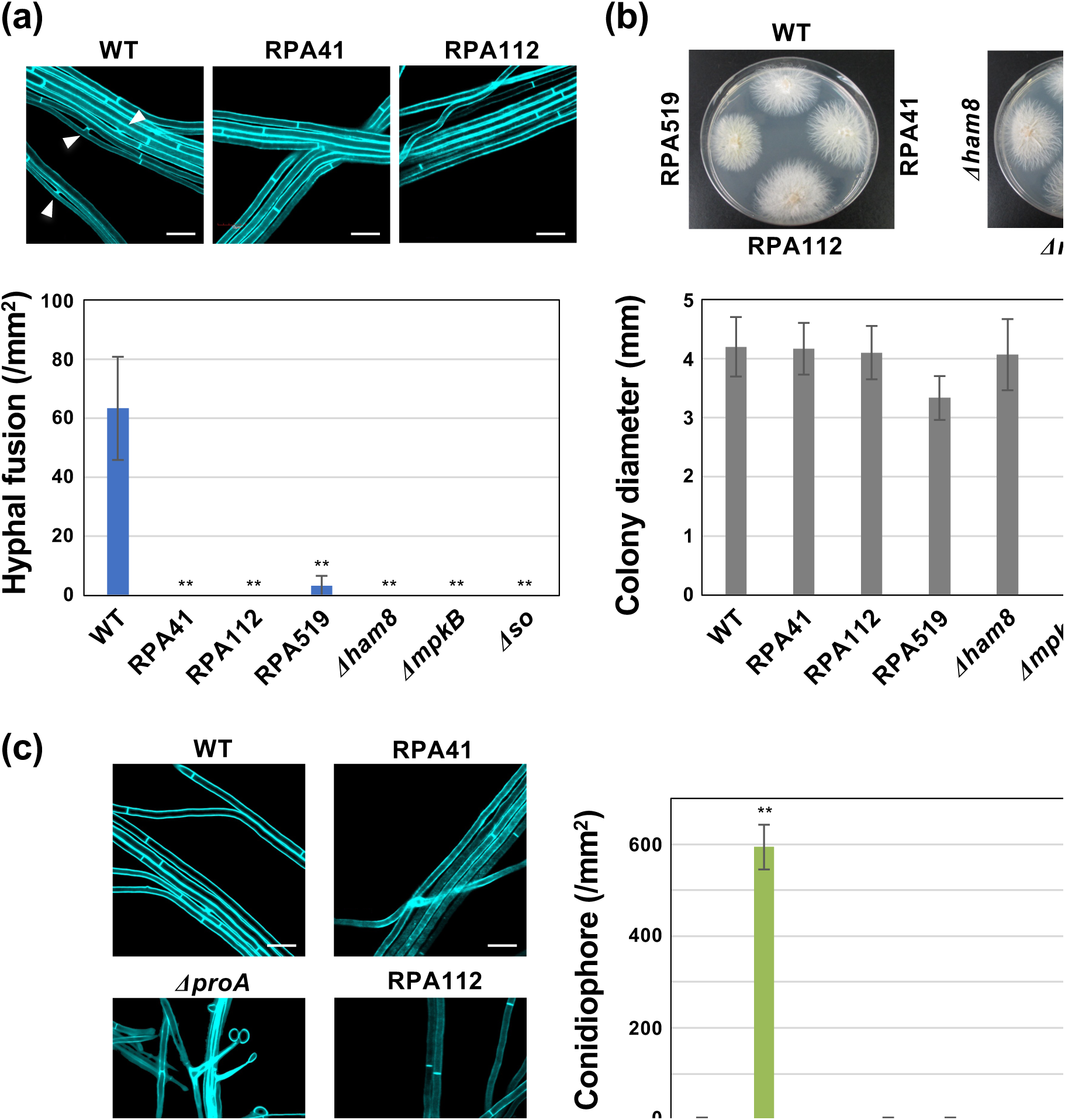
Isolation and characterization of hyphal fusion-deficient mutants of *Epichloë festucae.* **(a)** Hyphal fusion of *E. festucae* wild type (WT), strains RPA41, RPA112, RPA519, *ham8* mutant (*Δham8*), *mpkB* mutant (*ΔmpkB*) and *so* mutant (*Δso*) on water agar. Arrowheads indicate hyphal fusion events. The number of hyphal fusions was counted using a fluorescence microscope after staining with Calcofluor white. Data represent means ± standard error from 6 sites from three colonies of each strain. Data marked with asterisks are significantly different from WT as assessed by two-tailed Student’s *t* tests: \*\**P* < 0.01. Bars = 10 µm. **(b)** Colony morphology and growth of *E. festucae* WT and strains RPA41, RPA112, RPA519, *Δham8, ΔmpkB* and *Δso* on PDA after 14 days of cultivation. Colony diameters represent means ± standard error from three colonies of each strain. Two-tailed Student’s *t* tests showed no significant difference compared to the WT strain. **(c)** Conidiophore formation of *E. festucae* WT, *ΔproA,* RPA41, RPA112, RPA519, *Δham8, ΔmpkB* and *Δso* on water agar. The number of conidiophores was counted using a fluorescence microscope after staining with Calcofluor white. Data represent means ± standard error from 12 sites from three colonies of each strain. Data marked with asterisks are significantly different from WT as assessed by two-tailed Student’s *t* tests: \*\**P* < 0.01. Bars = 10 µm.

### 2.2 Analysis of Growth and Spore Formation in REMI Mutants

Growth on PDA medium was compared among the wild-type strain, the three REMI mutant strains isolated in the screening, and the *Δso*, *ΔmpkB*, and *Δham8* knockout strains. Colony morphology and growth rate were evaluated after two weeks of cultivation. Overall colony growth of the REMI mutants was comparable to that of the wild type, although RPA519 showed a slightly reduced growth rate. A moderate increase in aerial mycelium was observed in the *Δso* strain and, to a lesser extent, in RPA112 and RPA519. No other marked differences in colony morphology were detected among the tested strains during cultivation on PDA medium (Figure 1b).

Conidiation, which is rarely observed in wild-type *E. festucae* grown on PDA medium, has been reported to increase significantly in several previously characterized hyphal fusion-deficient strains such as *ΔnoxA* and *ΔproA* (Kayano et al., 2013; Tanaka et al., 2013). To assess whether this phenotype is shared by the newly identified mutants, conidial production was analyzed in the three REMI mutants and in the *Δso*, *ΔmpkB*, and *Δham8* strains. No increase in conidial production was detected in any of these mutants or knockout strains (Figure 1c). These findings indicate that the genes disrupted in RPA41, RPA112, and RPA519, as well as *so*, *mpkB*, and *ham8*, are essential for hyphal fusion but do not contribute to the regulation of conidiogenesis.

### 2.3 Analysis of Symbiotic Infection Ability of Hyphal Fusion-Deficient Mutants

To evaluate their ability to establish symbiotic infection, the three hyphal fusion-deficient mutants were inoculated into perennial ryegrass. For comparison, the *Δso* mutant, which has been reported to cause severe stunting of host plants, was included in the assay. Among the inoculated plants, those infected with RPA112 and RPA519 exhibited growth inhibition to a similar extent as that observed in the *Δso* mutant, often leading to pronounced stunting and eventual plant death (Figure 2a). In contrast, no infected plants were obtained when plant were inoculated with strain RPA41, a phenotype similar to that of the *ΔmpkB* mutant (Tanaka et al., 2020). These results indicate that all three hyphal fusion-deficient mutant strains isolated in this study exhibit defects in establishing a normal symbiotic interaction with the host plant.

**FIGURE 2.**
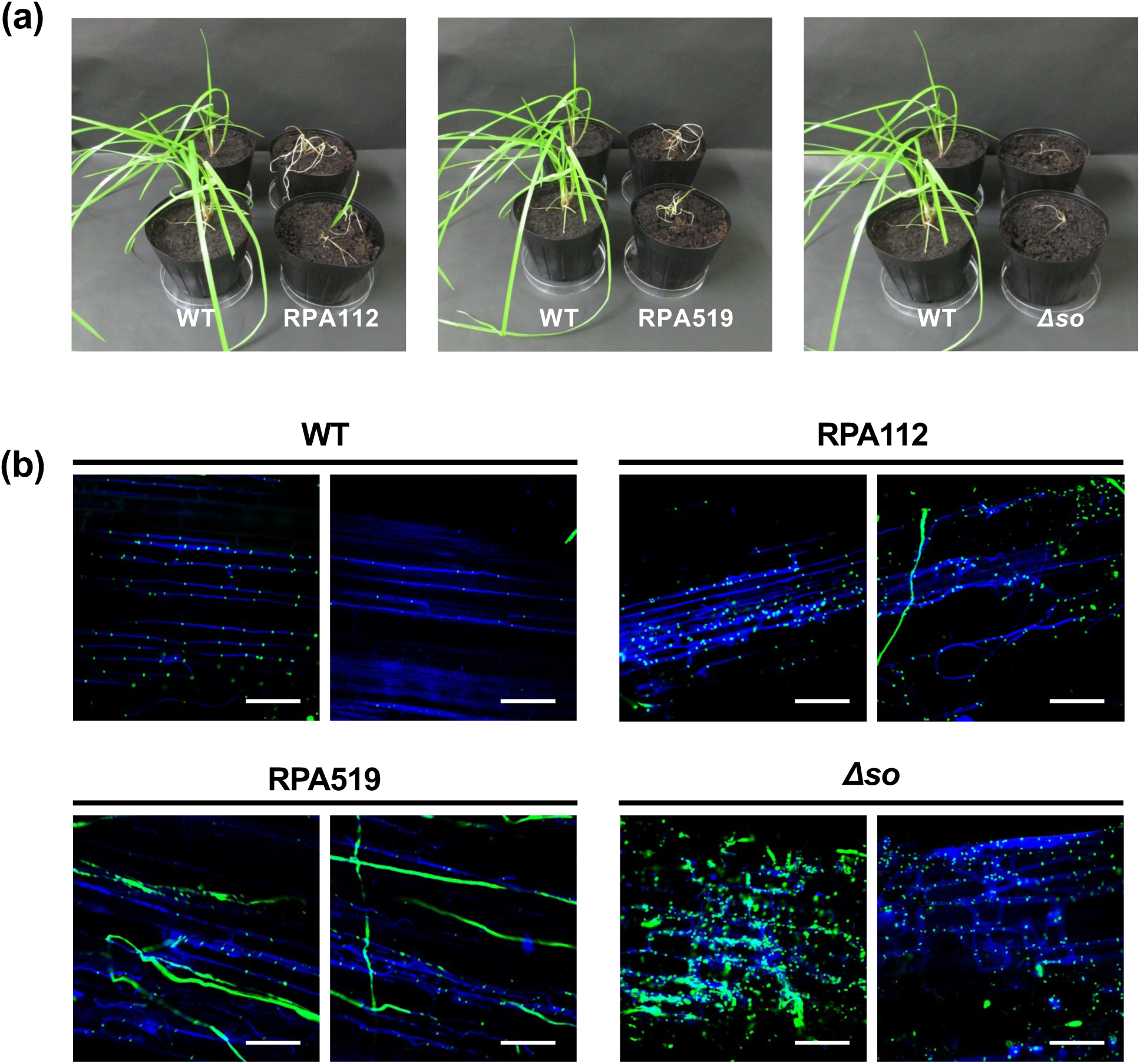
Plant infection phenotypes in *Epichloë festucae* hyphal fusion-deficient mutants. **(a)** Phenotypes of perennial ryegrass plants infected with *E. festucae* wild type (WT), strains RPA519 and RPA112, or *Δso*. Photographs were taken 8 weeks after inoculation. **(b)** Hyphal colonization patterns in infected pseudostem tissues stained with aniline blue and WGA-Alexa Fluor 488, and observed by confocal laser scanning microscopy. Bars = 50 µm.

The hyphae of strains RPA112, RPA519, and the *Δso* strain colonizing host plants were stained with aniline blue and WGA-Alexa Fluor 488 to observe hyphal growth and septation within host tissues. In contrast to the wild-type strain, where hyphae extended regularly within intercellular spaces, strains RPA112 and RPA519, as well as the *Δso* mutant, exhibited abnormal characteristics such as multiple hyphae extending into a single intercellular space, irregular septation, and hyphae extending beyond intercellular boundaries (Figure 2b). These observations indicate that RPA112 and RPA519 have lost the ability to maintain coordinated symbiotic growth with the host plant.

### 2.4 Identification of Genes Disrupted in Hyphal Fusion-Deficient Mutants

To identify the genes disrupted in the hyphal fusion-deficient mutants, the insertion patterns of the pNPP1 vector in the genomes of strains RPA41, RPA112, and RPA519 were examined (Figure 3). Total genomic DNA from each mutant was digested with *EcoR*I, *Cla*I (each containing a single restriction site within the vector), or *Eco*RV (which lacks a site in the vector), followed by Southern blot hybridization using the pNPP1 vector as a probe. A single hybridization band was detected in the *Eco*RV-digested genomic DNA of all three mutants, suggesting the presence of a single vector insertion site. Digestion with *Eco*RI and *Cla*I yielded a band of approximately 6 kb in strains RPA41 and RPA519, corresponding to the full vector length, indicating multiple tandem insertions of the vector at the same genomic locus. Based on the sizes of the hybridizing bands, genomic DNA digested with *Eco*RI or *Cla*I was separated by electrophoresis, the relevant fragments were excised from the gel, and self-ligated. The regions flanking the insertion sites were then amplified by inverse PCR with the ligated DNA using the primer set PtrpC-2/hph-2, and the amplified DNA fragments were sequenced. From the genomic sequences adjacent to the pNPP1 vector, the insertion sites of each REMI transformant were determined.

**FIGURE 3.**
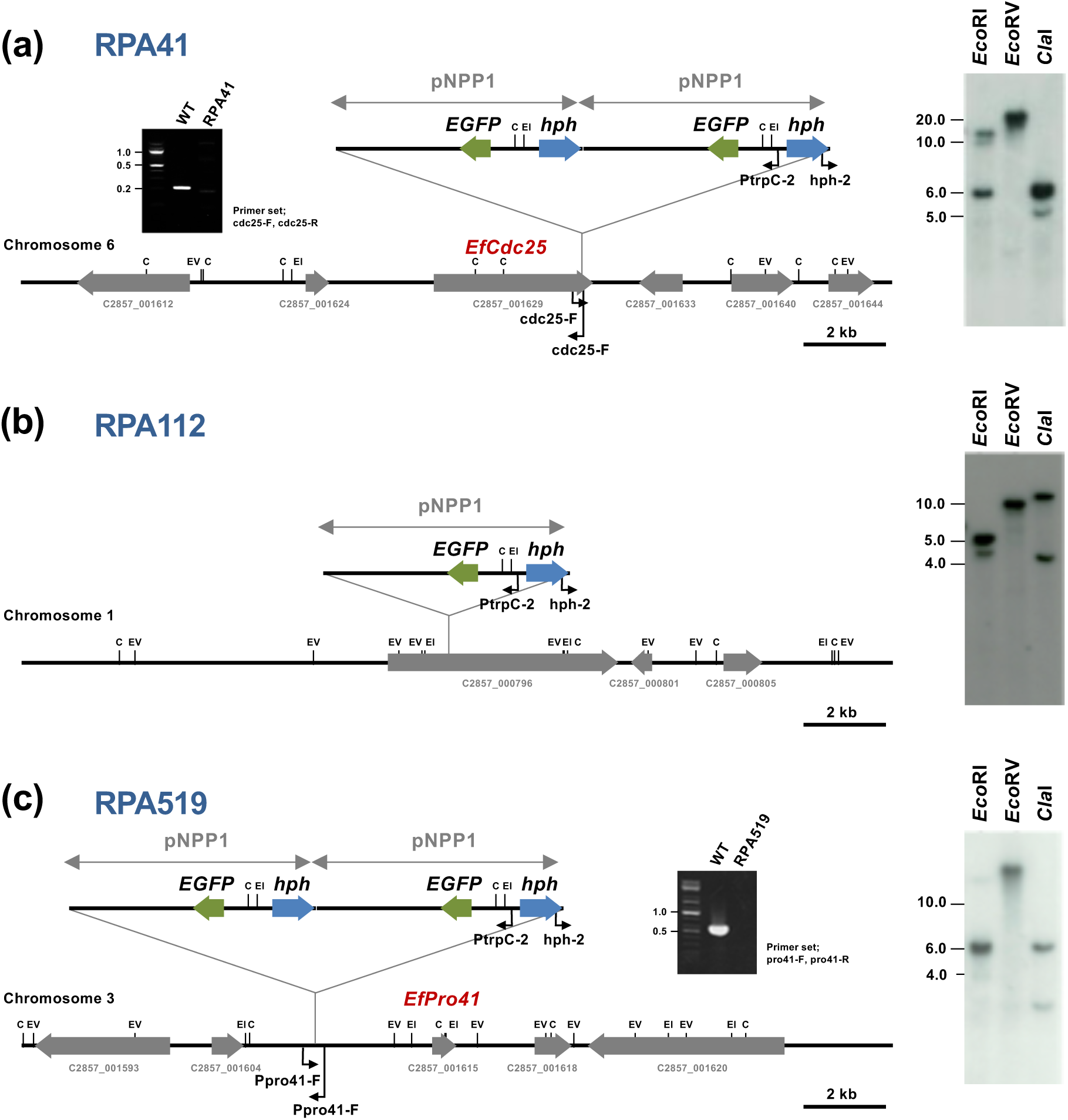
Identification of genes disrupted in *Epichloë festucae* hyphal fusion-deficient mutants. (Left) Physical maps of pNPP1-tagged locus in RPA41 **(a)**, RPA112 **(b)** and RPA519 **(c)**, showing the plasmid insertion site in the mutants. (Right) Southern blot analysis of genomic DNA from hyphal fusion-deficient mutants. Genomic DNA (0.2 µg) was digested with *Eco*RI, *Eco*RV, or *Cla*I, separated by electrophoresis in 0.8% agarose gel, and hybridized with [^32^P]-labelled pNPP1. (Insets) PCR amplification of insertion sites of RPA41 and RPA519 using genomic DNA from wild type and REMI mutants.

For RPA41, the vector insertion site was located within the coding region of the *cdc25* gene (locus tag C2857_001629) on chromosome 6, which encodes a guanine nucleotide exchange factor (GEF) for the small GTPase Ras. PCR using primers for the *cdc25* gene showed that the approximately 220 bp band detected in the wild-type strain was absent in RPA41, confirming vector insertion into the predicted *cdc25* locus (Figure 3a).

For RPA112, the vector insertion site was predicted to be located in the C2857_000796 gene on chromosome 1, encoding a hypothetical protein (Figure 3b). A disruption mutant of C2857_000796 was generated; however, no defect in hyphal fusion was observed (data not shown). It is possible that, during REMI transformation, genomic deletions occurred outside the vector insertion site. Therefore, further analyses such as whole-genome sequencing, will be required in future studies to identify the gene responsible for the observed phenotype of RPA112.

For RPA519, the vector insertion site was predicted to be located approximately 2.8 kb upstream of the *pro41* gene (C2857_001615) on chromosome 3 (Figure 3c). Pro41 is known to be involved in sexual reproduction in *Sordaria macrospora* and hyphal cell fusion in *N. crassa* (Nowrousian et al., 2007; Fu et al., 2011). Pro41 has been proposed as a component of the fungal NADPH oxidase complex (Takemoto and Scott, 2023), and we have previously reported that Pro41 is essential for hyphal fusion and symbiotic infection in *E. festucae* (Tanaka et al., 2020). The partial hyphal fusion observed in RPA519 (Figure 1a) is presumed to result from disruption of the promoter region rather than the coding region of *pro41* (Figure 3c). In the subsequent analyses, we focused on the functional characterization of *cdc25*, which was newly identified as an essential factor for hyphal fusion and the establishment of symbiosis in *E. festucae*.

### 2.5 Characterization of *E. festucae cdc25* Knockout Strains

The *E. festucae* Cdc25 protein (EfCdc25; C2857_001629) consists of 1,160 amino acids and shares conserved structural features typical of fungal Ras guanine nucleotide exchange factors (RasGEFs) (Figure S1). The N-terminal region contains an Src homology 3 (SH3) domain with a peptide ligand-binding site, which is generally involved in protein–protein interactions and, in *S. cerevisiae* Cdc25, mediates binding to adenylate cyclase (Mintzer and Field, 1999). The central region is compositionally rich in serine and threonine residues, a characteristic feature of intrinsically disordered regulatory regions in fungal signaling proteins (van der Lee et al., 2014). The C-terminal half contains a Ras exchange motif (REM) domain, a RasGEF catalytic domain, and a conserved Ras-binding site, all essential for Ras interaction and activation. These structural features indicate that EfCdc25 functions as a canonical Ras-specific GEF, likely mediating nucleotide exchange through direct Ras binding (Figure S1).

*cdc25* knockout strains were generated using homologous recombination. Gene disruption vectors, pNPP233 (targeting the entire *cdc25* gene) and pNPP234 (targeting the Ras-binding domain of *cdc25*) were constructed (Figure 4a). One *Δcdc25* strain (Δcdc25-13) and five *Δcdc25-RasBD* strains (*Δcdc25-RasBD*-16, -17, -18, -25, and -26) were obtained. Southern hybridization confirmed that all candidates carried the predicted integration of the knockout vectors (Figure 4b). Both *Δcdc25* and *Δcdc25-RasBD* strains showed no significant abnormality in colony growth; however, no hyphal fusion was observed (Figure 4c), confirming that *cdc25* is required for hyphal fusion in *E. festucae*.

**FIGURE 4.**
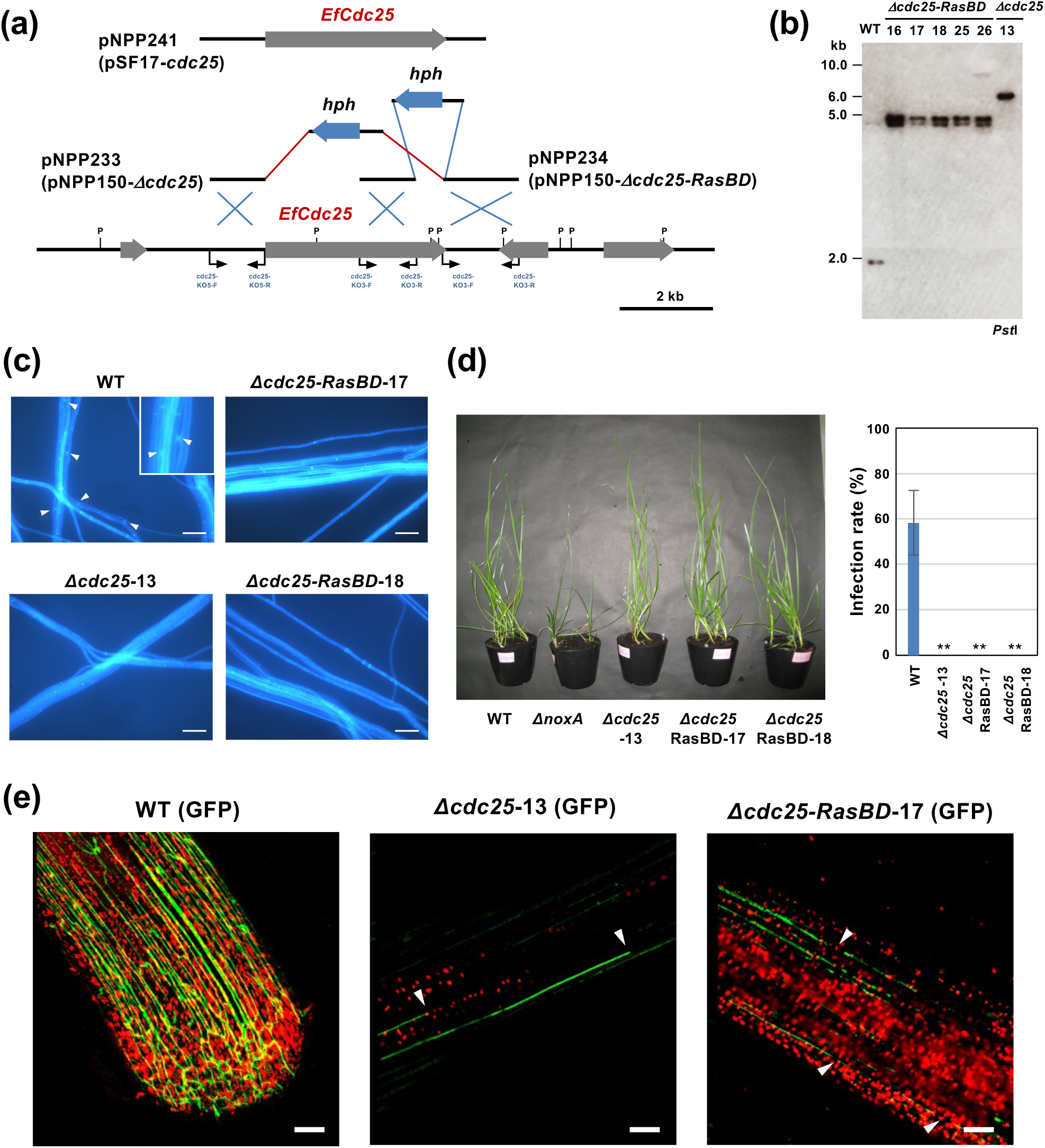
Characterization of *Epichloë festucae cdc25* knockout strains. **(a)** Physical map of the *cdc25* wild-type genomic region and linear insert of *cdc25* replacement constructs pNPP233 (pNPP150-*Δ*cdc25), pNPP234 (pNPP150-*Δcdc25-RasBD*) and complementation construct pNPP241 (pSF17-cdc25). P, *Pst*I. **(b)** Autoradiograph of Southern blot of *Pst*I genomic digests of *E. festucae* wild-type (WT), *Δcdc25* and *Δcdc25-RasBD* strains hybridized with [^32^P]-labeled pNPP150-*Δcdc25*. The sizes of marker DNA fragments are given in kb. **(c)** Hyphal fusion of *E. festucae* WT, *Δcdc25* and *Δcdc25-RasBD* strains on water agar. Hyphal fusion events were observed using a fluorescence microscope after staining with Calcofluor white. Bars = 100 µm. **(d)** Phenotypes of perennial ryegrass infected with *E. festucae* WT, *ΔnoxA*, and *cdc25* knockout strains. Infection rates were assessed 8 weeks after inoculation. Data represent means ± standard error from 3 independent experiments. Data marked with asterisks are significantly different from WT as assessed by two-tailed Student’s *t* tests: **P < 0.01. **(e)** Hyphal growth of GFP-labeled *E. festucae* in host plant. Seedlings of perennial ryegrass were inoculated with *E. festucae* wild type, *Δcdc25*-13 or *Δcdc25-RasBD*-17 transformed with a GFP expression vector. Photographs were taken 10 days after inoculation. Arrowheads indicate fragmentation of *E. festucae* hyphae. Bars = 50 μm.

Strains *Δcdc25-13*, *Δcdc25-RasBD-17*, and *Δcdc25-RasBD-18* were inoculated into perennial ryegrass to evaluate their ability to establish symbiotic infection. While infection by the wild-type strain was confirmed, no infection was detected in host plants inoculated with either *Δcdc25* or *Δcdc25-RasBD* strains, showing phenotype similar to that of strain RPA41. These observations demonstrate that EfCdc25 is essential for symbiotic colonization of *E. festucae* in its host plant (Figure 4d). For complementation, a genomic fragment containing approximately 1.6 kb upstream and 1.1 kb downstream of the *cdc25* gene was introduced into the *Δcdc25* and *Δcdc25-RasBD* strains. The resulting complemented strains regained both hyphal fusion ability and symbiotic infection, further confirming that EfCdc25 mediates hyphal fusion and symbiotic development in *E. festucae* (Figure S2).

To determine the stage of infection at which the *cdc25* mutants lost symbiotic ability, GFP-expressing *Δcdc25* and *Δcdc25-RasBD* strains were generated, and their growth within host tissues was examined. After inoculation of the leaf-sheath base of perennial ryegrass seedlings, initial infection by both knockout strains was confirmed; however, fragmented hyphae and reduced fungal biomass were consistently observed, in contrast to the well-developed hyphal networks formed by the wild-type strain (Figure 4e). In mature plants, hyphae of either *Δcdc25* or *Δcdc25-RasBD* strains were not detected, indicating that loss of *cdc25* function prevents synchronized hyphal extension with host growth during symbiotic colonization by the endophyte.

### 2.6 Analysis of the Interaction between Cdc25 and Ras-type Small GTPases

*E. festucae* Cdc25 exhibits a domain organization typical of fungal Ras guanine nucleotide exchange factors (Figure S1). In the *E. festucae* genome, five Ras-type small GTPases, RasA, RasB, RasC, KrevA, and RhbA, were identified (Figure 5a; Kayano et al., 2018). To examine whether Cdc25 interacts with these Ras proteins, yeast two-hybrid assays were performed. Because cysteine residues near the C-terminus serve as geranylgeranylation sites for plasma membrane localization and may interfere with the assay, cysteine-to-alanine substitutions were introduced into each Ras protein. Among the five Ras family members, only RasB showed a clear interaction with Cdc25 (Figure 5b), suggesting that EfCdc25 functions as a RasB-specific guanine nucleotide exchange factor.

**FIGURE 5.**
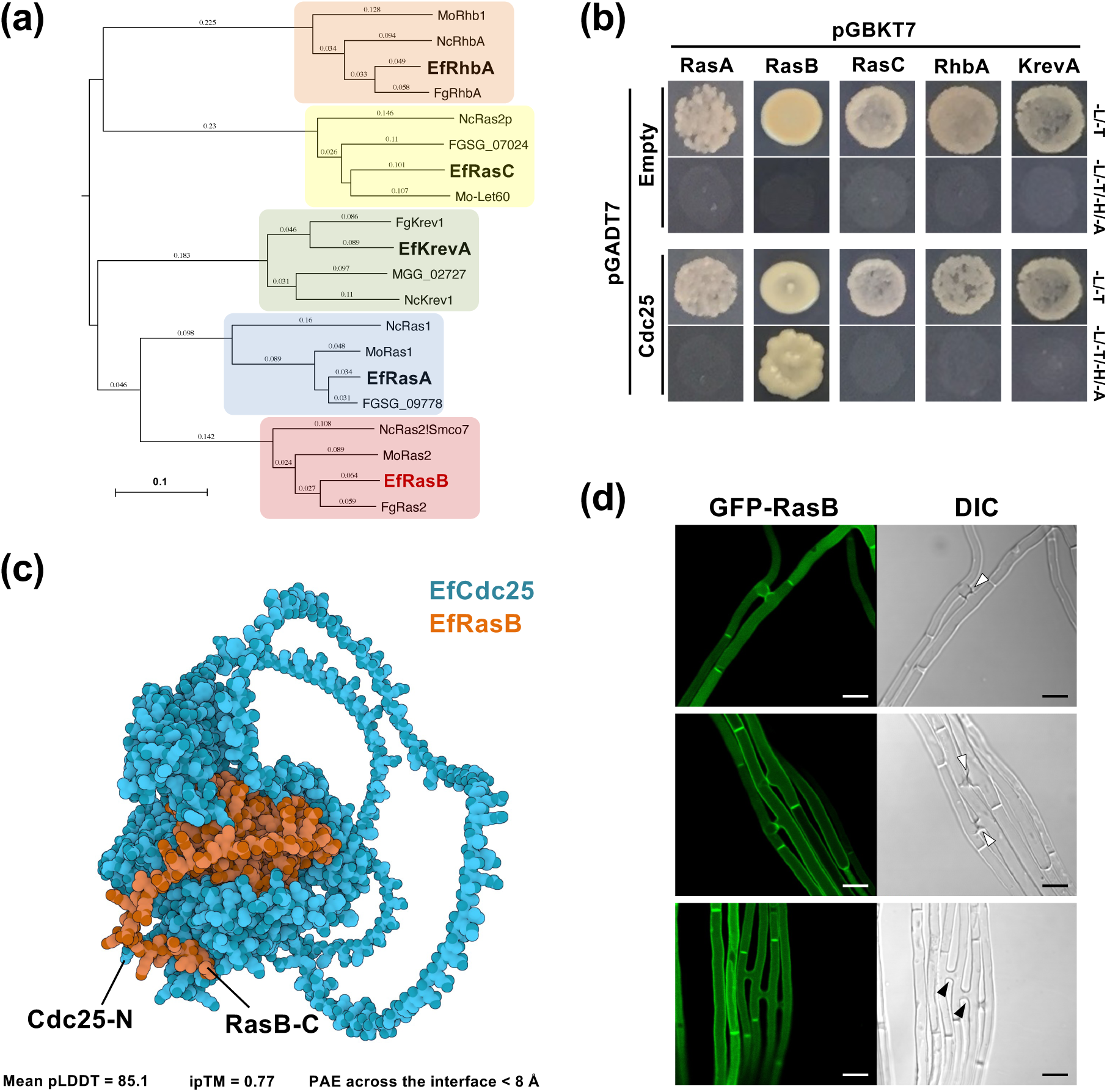
(a) Phylogenetic analysis of Ras GTPase of Ascomycota fungi. The tree was generated by the neighbor-joining method based on pairwise amino acid distances. The scale bar represents 0.1 amino acid substitutions per site. Ef, *Epichloë festucae*; Fg, *Fusarium graminearum*; Mo, *Magnaporthe oryzae*; Nc, *Neurospora crassa*. **(b)** Yeast two-hybrid assays for interactions between *E. festucae* Cdc25 and 5 Ras small GTPases. RasA, RasB, RasC, RhbA and KrevA have mutation in C-terminal plasma membrane localization signal. Yeast strain AH109 was transformed with prey and bait vectors, pGADT7 and pGBKT7, as indicated and plated on to SD medium lacking leucine and tryptophan (-L/-T) or lacking leucine, tryptophan, histidine and adenine (-L/-T/-H/-A). Growth on the latter indicates an interaction between bait and prey. **(c)** Predicted structure of the *E. festucae* Cdc25-RasB complex generated by AlphaFold3. The top-ranked model showed a mean pLDDT of 85.1, an interface predicted TM (ipTM) score of 0.77, and a predicted TM (pTM) score of 0.51. The predicted aligned error (PAE) across the interface was < 8 Å, indicating a stable and specific interaction between the RasGEF catalytic domain of Cdc25 and the Switch regions of RasB (Table S1). **(d)** Subcellular localization of GFP-RasB in hyphae of *E. festucae*. GFP-RasB was expressed under the control of a *TEF* promoter. Localization of GFP-RasB was observed for three independent transformants. White and black arrowheads indicate hyphal tips undergoing fusion and completed fusion sites, respectively. Bar = 5 μm.

*E. festucae* RasB possesses all conserved motifs characteristic of Ras-type small GTPases, including the P-loop, Switch I and II regions, and the C-terminal CaaX motif required for membrane anchoring via geranylgeranylation (Figure S3). To gain further insight into the EfCdc25-RasB association, a complex structure was modeled using AlphaFold3. The predicted model revealed a stable interface between the RasGEF catalytic domain of EfCdc25 and the Switch regions of EfRasB, with an interface predicted TM-score (ipTM) of 0.77 and a predicted TM-score (pTM) of 0.51. The ipTM reflects the confidence of the inter-chain interface, whereas the pTM represents the overall structural reliability of the complex. The average per-residue confidence (pLDDT) was approximately 85, and the mean predicted aligned error (PAE) across the interface was below 8 Å, indicating a high-confidence structural model. These computational analyses support a direct and specific interaction between EfCdc25 and EfRasB, consistent with the results of the yeast two-hybrid assay (Figure 5c, Table S1).

### 2.7 Cellular Localization of GFP-RasB and GFP-Cdc25

To investigate the intracellular localization of RasB and Cdc25 in *E. festucae* hyphae, transformants expressing GFP fusion proteins were generated. In strains expressing GFP-RasB, fluorescence was observed along the hyphal plasma membrane, with strong accumulation at the septa and at the apical region of actively growing hyphae immediately before fusion (Figure 5d). This polarized localization was transient and disappeared after fusion, suggesting that RasB accumulates at septa specifically during the pre-fusion stage. In contrast, in strains expressing GFP-Cdc25, weak GFP fluorescence was detected without a clear subcellular localization pattern. However, slightly intensified GFP patches were occasionally observed, suggesting a possible transient accumulation of EfCdc25 at specific sites preceding hyphal fusion, although further analysis is required to confirm this observation (Figure S4).

### 2.8 Attempted Disruption of *rasB* Gene in *E. festucae*

To determine whether RasB is required for hyphal fusion and symbiotic infection, as observed for Cdc25, disruption of the *rasB* gene was attempted. Gene replacement constructs containing genomic fragments flanking the *rasB* coding region were generated (Figure S5). Two types of disruption vectors were introduced into *E. festucae* protoplasts; however, no transformants carrying a disrupted *rasB* allele were obtained despite multiple independent transformation attempts. These results suggest that *rasB* is an essential gene in *E. festucae*, and its loss may cause severe growth defects or lethality. A similar essentiality of Ras-type GTPases has been reported in other filamentous fungi, including *Botrytis cinerea* (*rasA*; Li et al., 2023), and the rice blast fungus *Pyricularia oryzae* (syn. *Magnaporthe oryzae*) (*ras2*; Park et al., 2006).

### 2.9 Complementation of *Δcdc25* Strains with Constitutively Active RasB

Because Cdc25 is presumed to activate RasB by promoting its conversion to the GTP-bound form, the loss of hyphal fusion and symbiosis establishment observed in the *Δcdc25* strains was expected to result from the absence of RasB activation. To test this possibility, a constitutively active variant of RasB (CA-RasB), carrying a G16V substitution that inhibits GTP hydrolysis (Figure S3), was expressed in the *Δcdc25* strain. The resulting transformants were first examined for their ability to undergo hyphal fusion. Expression of CA-RasB in the *Δcdc25* restored hyphal fusion to a level comparable to that of the wild type (Figure 6a).

**FIGURE 6.**
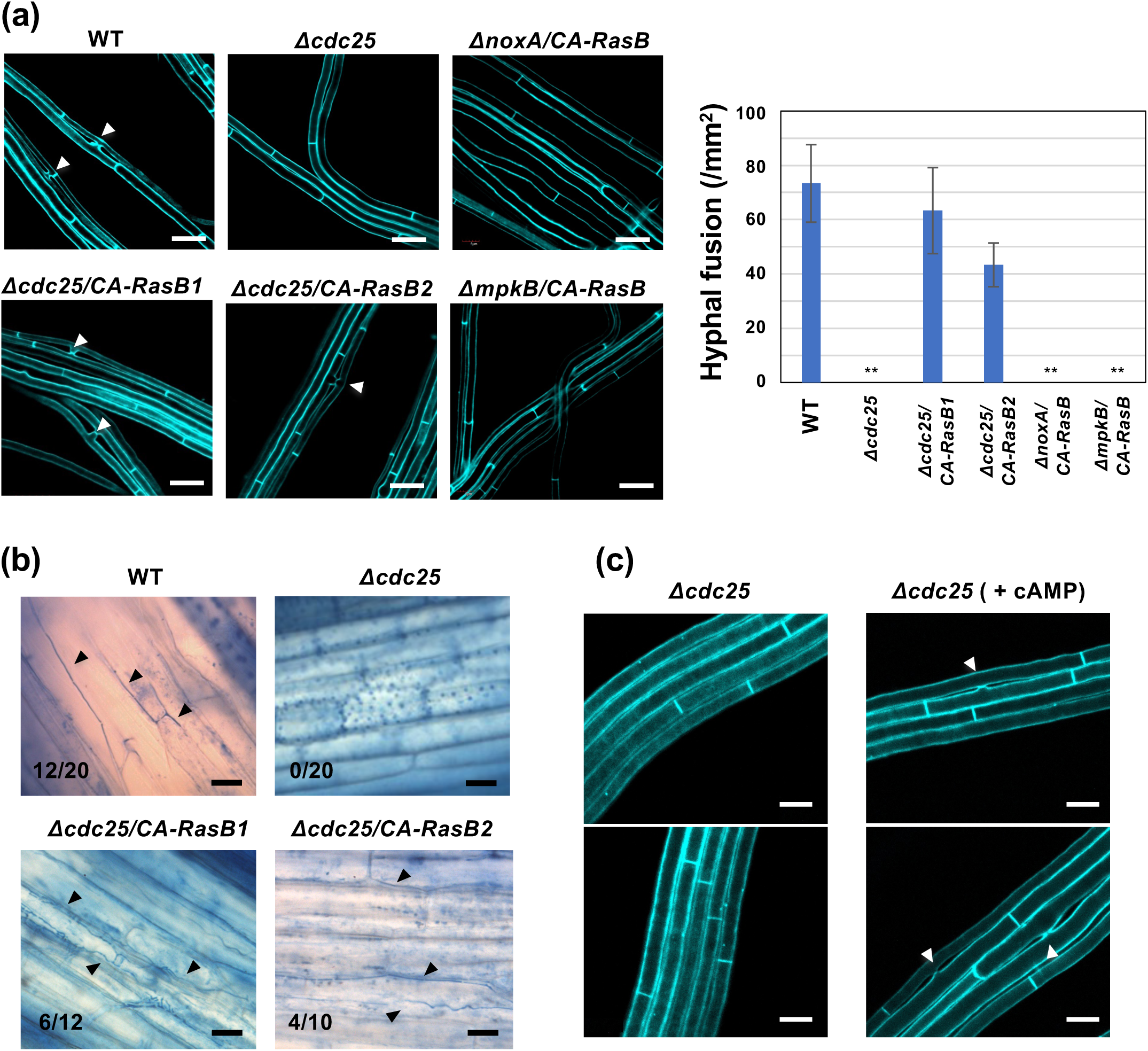
Complementation of *Epichloë festucae Δcdc25* strains with constitutively active RasB. **(a)** Hyphal fusion of *E. festucae* wild type (WT), *Δcdc25-13*, *Δcdc25-13* expressing constitutively active RasB (CA-RasB)*, ΔnoxA* or *ΔmpkB* expressing CA-RasB grown on water agar. The number of hyphal fusions was counted using fluorescence microscopy after staining with Calcofluor white. Data represent means ± standard error from six observation sites across three colonies per strain. Asterisks indicate significant differences from WT as assessed by two-tailed Student’s *t* tests: **P < 0.01. Bars = 10 µm. **(b)** Colonization of *E. festucae* strains in host plant perennial ryegrass. Pseudostem of perennial ryegrass was inoculated with each strain, stained with aniline blue, and observed under light microscopy. Infection rate (number of infected/inoculated plants) is shown in each panel. Bars = 30 µm. **(c)** Partial recovery of hyphal fusion in *E. festucae Δcdc25* strain on water agar supplemented with 10 mM cyclic AMP (cAMP). Bars = 5 µm.

To further determine whether RasB activation could also restore symbiotic infection, wild-type, *Δcdc25*, and *Δcdc25 /CA-RasB* strains were inoculated into perennial ryegrass, and fungal colonization was assessed. Hyphal growth within host tissues was evident in plants infected with the wild-type strain but absent in those inoculated with *Δcdc25*. In contrast, plants inoculated with the *Δcdc25*/CA-RasB strain displayed restored endophytic colonization comparable to that of the wild type (Figure 6b). These results demonstrate that constitutive activation of RasB is sufficient to rescue both hyphal fusion and symbiotic infection in the absence of Cdc25, indicating that Cdc25 functions upstream of RasB in the signaling pathway regulating hyphal fusion and symbiotic development in *E. festucae*.

Interestingly, when CA-RasB was overexpressed in the wild-type background, abnormal hyphal swelling was observed (Figure S6). This morphological abnormality was not prominent in the *Δcdc25* strain, suggesting that proper regulation of active RasB levels is required for maintaining normal hyphal morphology.

### 2.10 Constitutively Active RasB Does Not Restore Hyphal Fusion in the *ΔmpkB* and *ΔnoxA* Strains

The MAP kinase MpkB plays a central role in hyphal fusion and the establishment of symbiosis, and *ΔmpkB* mutants exhibit phenotypes closely resembling those of *Δcdc25* strains, including a complete loss of hyphal fusion and an inability to infect host plants (Tanaka et al., 2020). To clarify the relationship between RasB signaling and the MAP kinase pathway, CA-RasB was expressed in the *ΔmpkB* background, but CA-RasB failed to restore hyphal fusion ability in the *ΔmpkB* strain. These results indicate that, as previously reported in other fungi, the Cdc25-RasB module functions upstream of MpkB.

NADPH oxidase NoxA and its regulatory components are required for hyphal fusion and normal host infection in *E. festucae*, but their roles in the signaling pathway remain unclear (Takemoto et al., 2006b). Expression of CA-RasB in the *ΔnoxA* mutant did not complement the fusion defect (Figure 6a). These observations suggest that NoxA may act downstream of the Cdc25-RasB module or function in a parallel pathway.

### 2.11 Partial Restoration of Hyphal Fusion in the *Δcdc25* Strain by Exogenous cAMP

In the signaling pathway involving Cdc25 and RasB, adenylate cyclase, the enzyme responsible for cyclic AMP (cAMP) synthesis, is presumed to be activated downstream. Cdc25 contains an N-terminal SH3 domain (Figure S1), which in *Saccharomyces cerevisiae* mediates binding to adenylate cyclase (Mintzer and Field, 1999). To examine whether exogenous cAMP could compensate for the loss of Cdc25, the *Δcdc25* strain was cultured on water agar medium supplemented with 10 mM cAMP, and hyphal fusion was assessed. Although hyphal fusion was completely absent in the *Δcdc25* strain, the addition of cAMP partially restored hyphal fusion (Figure 6c). These results indicate that the Cdc25-RasB module promotes hyphal fusion, at least in part, through a cAMP-dependent signaling pathway.

## 3 DISCUSSION

### 3.1 Signaling Coordination Underlying Symbiotic Steady State and Hyphal Fusion in *Epichloë* Endophytes

The establishment of a symbiotic association between *E. festucae* and perennial ryegrass requires the fungus to maintain a finely tuned balance between proliferation and restraint within host tissues. Hyphal fusion is presumed to be a central process underlying this steady state of symbiotic growth, allowing the formation of a continuous hyphal network that facilitates intercellular communication and synchronized growth with the host plant. Previous studies have revealed that mutants defective in symbiosis also lack the ability to undergo hyphal fusion, indicating that these two processes are tightly linked (Kayano et al., 2013; Tanaka et al., 2020). Disruption of genes encoding NADPH oxidase components NoxA, NoxR, and RacA; MAP kinase MpkA; the STRIPAK complex component MobC; the scaffold protein SO; or transcriptional regulators such as ProA and NsiA results in nearly identical phenotypes—loss or severe reduction of hyphal fusion and uncoordinated hyphal proliferation *in planta* (Takemoto et al., 2006a; Tanaka et al., 2006, 2008, 2013, 2020; Charlton et al., 2012; Becker et al., 2015; Green et al., 2017). Interestingly, most fusion-defective mutants cause pronounced host stunting, whereas others, such as *ΔmpkB*, *Δste12*, *Δcdc24*, and *Δcdc25*, exhibit a complete or partial loss of infectivity and systemic colonization in the host plant (Kayano et al., 2018; Tanaka et al., 2020; Figure 4). These observations suggest that distinct signaling thresholds and partially independent pathways operate at different stages of symbiotic establishment, ensuring that hyphal behavior remains coordinated and compatible with host growth patterns.

### 3.2 Evolutionary Rewiring of the Conserved Cdc25-Ras Signaling Module from Pathogenicity to Mutualism

In several filamentous fungi, particularly plant pathogens, Cdc25 has been identified as a Ras-specific guanine nucleotide exchange factor (RasGEF) that governs infection-related morphogenesis and virulence. In *Fusarium graminearum*, deletion of *FgCdc25* (the sole RasGEF in this species) causes severe defects in vegetative growth, sexual reproduction, and pathogenicity, phenotypes that can be partially alleviated by exogenous cAMP. FgCdc25 specifically activates Ras2, thereby linking the Ras-cAMP/PKA pathway to MAPK cascades that regulate hyphal differentiation, trichothecene biosynthesis, and host invasion (Chen et al., 2020). Similar functions have been described in *Colletotrichum higginsianum*, where deletion of *ChCdc25* abolishes conidiation and appressorium formation, leading to complete loss of pathogenicity (Yan et al., 2020). In *Ustilago maydis*, the Cdc25 homolog Sql2 activates Ras2 and is indispensable for the transition from the yeast-like to the filamentous growth required for infection (Müller et al., 2003). Collectively, these studies underscore the conserved role of Cdc25-mediated Ras activation as a central regulatory node coupling morphogenesis and virulence in diverse fungal pathogens.

In contrast, *E. festucae* appears to have evolutionarily repurposed this ancient Ras regulatory module for mutualistic symbiosis rather than pathogenic invasion. While the overall Ras signaling architecture is conserved, its physiological output is fundamentally different. Instead of triggering invasive structures such as appressoria or penetration hyphae, Cdc25-mediated RasB activation in *E. festucae* regulates hyphal fusion and coordinated growth within the intercellular spaces of the host. This shift in signaling outcome likely reflects an evolutionary adaptation in which an ancestral virulence-associated pathway was co-opted to maintain communication and compatibility with the host plant. Such functional reconfiguration mirrors the broader principle that mutualistic fungi often retain pathogenicity-related genes and signaling networks but employ them for controlled, non-destructive interaction with the host.

### 3.3 Cross-Talk and Coordination among Cdc25-RasB, cAMP, MAPK, and Redox Pathways for Symbiotic Homeostasis in *E. festucae*

A conceptual overview of the interconnected signaling pathways is illustrated in Figure 7. In pathogenic species such as *P. oryzae*, *C. higginsianum*, and *F. graminearum*, Ras-dependent activation of MAPKs is essential for infection structure formation, including appressoria and penetration hyphae (Park et al., 2006; Chen et al., 2020). In *E. festucae*, MpkB regulates both hyphal fusion and the establishment of symbiotic infection (Tanaka et al., 2020), and the inability of CA-RasB to complement *ΔmpkB* is consistent with MpkB acting as a non-redundant downstream requirement. At the same time, partial restoration of fusion in *Δcdc25* by exogenous cAMP suggests convergence of RasB signaling on the adenylate cyclase-PKA axis, which is compatible with the presence of an SH3 domain in EfCdc25 that can mediate binding to adenylate cyclase in fungi (Mintzer and Field, 1999). Phenotypes of *E. festucae acyA* mutants (slow growth, hyperbranching, and over colonization in host plant) further indicate that cAMP signaling only partially contributes to the hyphal growth synchronization with the host plant (Voisey et al., 2016). In parallel, the NADPH oxidase branch (NoxA, NoxR, RacA) provides redox cues that influence hyphal communication and polarity; loss of these components disrupts fusion and *in planta* organization (Tanaka et al., 2006; Takemoto et al., 2006, 2011), implying that ROS production furnishes a permissive context for MAPK-mediated controls.

**FIGURE 7.**
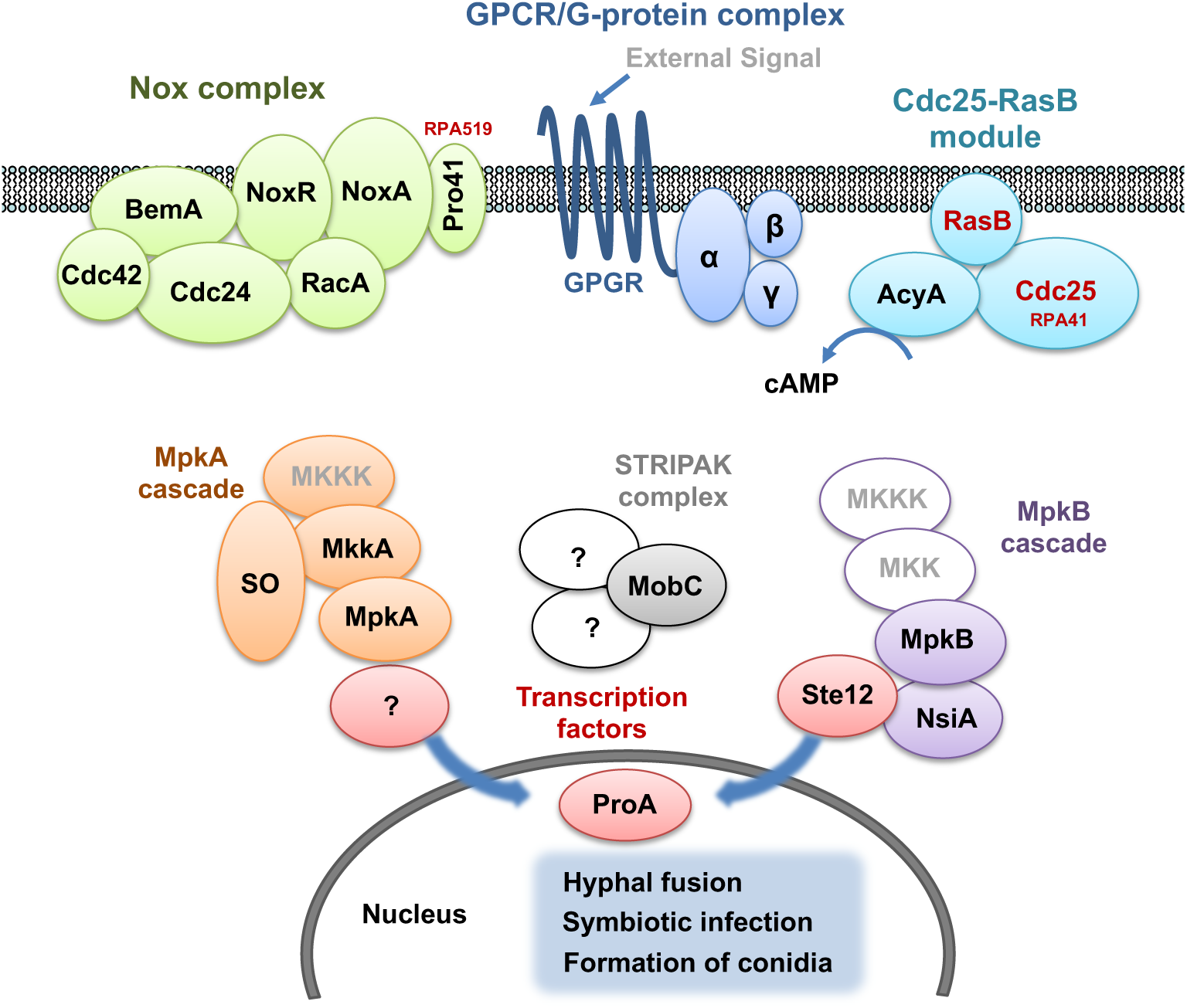
Schematic model showing coordinated regulation of hyphal fusion, symbiotic infection, and conidiation by the Cdc25–RasB module (cell division cycle 25–Ras GTPase), mitogen-activated protein kinase (MAPK) cascades, striatin-interacting phosphatase and kinase (STRIPAK) complex, and NADPH oxidase (Nox) complex in *Epichloë festucae*.

As summarized in Figure 7, the Cdc25-RasB arm, the Nox complex, and the MAPK and STRIPAK cascades operate in close coordination with extensive cross-talk. Scaffold and organizing proteins such as SO and MobC provide physical and functional coupling among modules that ultimately converge on transcriptional regulators including Ste12, NsiA, and ProA (Tanaka et al., 2013, 2020). Current models in filamentous fungi place Ras as an upstream hub that branches to a cAMP/PKA arm and a MAPK arm, with bidirectional tuning between PKA and MAPK depending on context (Turrà et al., 2014; Fortwendel, 2015). In pathogenic fungi, both arms drive invasion programs; in *E. festucae*, the same core circuitry is reconfigured to promote hyphal fusion, tip polarity, restraint of branching, and host-synchronized growth. Elucidating how these interconnected modules sense and integrate host-derived cues will be essential for understanding the molecular basis of fungal symbiotic homeostasis.

## 4 Experimental Procedures

### 4.1 Biological Materials, Growth Conditions, and Inoculation

*Epichloë festucae* strains (Table S2) were cultured on potato dextrose agar (PDA) or water agar (3% agar, w/v) at 23°C. Endophyte-free seedlings of perennial ryegrass (*Lolium perenne* cv. Yatsukaze) were inoculated with *E. festucae* following the method of Latch and Christensen (1985). Plants inoculated with the endophyte were grown as previously described (Tanaka et al., 2005).

### 4.2 DNA Preparation and Hybridization

Fungal genomic DNA was extracted from cultured mycelium as described previously (Byrd et al., 1990) or using an Extract-N-Amp Plant PCR Kit (Sigma-Aldrich, USA). Southern hybridization was performed following the procedure of Kayano et al. (2018).

### 4.3 Construction of Vectors

PCR amplification of genomic and plasmid DNA templates was performed using PrimeSTAR Max DNA polymerase (Takara Bio, Japan) or GoTaq Green Master Mix (Promega, USA). PCR fragments were cloned into plasmids using the In-Fusion HD Cloning Kit (Takara Bio). Vectors for gene knockout, complementation, and gene expression in *E. festucae* are listed in Table S3. Vectors for yeast two-hybrid assay are listed in Table S4. Sequences of primers used for the vector construction are listed in Table S5.

### 4.4 *E. festucae* Transformation and Restriction Enzyme-Mediated Integration Mutagenesis

Protoplasts of *E. festucae* were prepared according to Kuroyanagi et al. (2022). Transformation was performed with 5 µg of either circular plasmids or linearized plasmids (for gene knockout) using the same procedure. In this study, we generated a knockout mutant of the *so* and confirmed its previously reported phenotype (Figure S7; Charlton et al., 2012). For restriction enzyme-mediated integration (REMI) mutagenesis (Sánchez et al., 1998), protoplasts were transformed with 5 µg of *Pst*I-linearized pNPP1 (Kayano et al., 2013), together with 1 unit of *Pst*I added to the plasmid-protoplast mixture. Transformants were selected on PDA containing hygromycin (150 µg/ml), geneticin (400 µg/ml), or both antibiotics for co-transformation. Transformants expressing EGFP were identified under a BX51 fluorescence microscope (Olympus, Japan).

### 4.5 Screening of Hyphal Fusion-Deficient Mutants

REMI transformants grown on PDA supplemented with 150 µg/ml hygromycin were inoculated onto water agar (3% agar, w/v) and incubated at 23 °C for 1–2 weeks. Small agar blocks containing surface hyphae were excised, and 10% Calcofluor white solution (Sigma-Aldrich, USA) was applied directly to the agar surface for staining. The stained hyphae were examined under a BX51 fluorescence microscope (Olympus, Japan). Transformants showing no visible hyphal fusion were selected as candidate mutants. These candidates were re-examined using the same procedure, and three fusion-deficient mutants were finally isolated from approximately 1,200 REMI transformants.

### 4.6 Microscopy

Images of GFP-labelled or aniline blue/WGA-AF488-stained *E. festucae* in host tissues, GFP-tagged proteins in fungal hyphae, and Calcofluor white-stained hyphae (Sigma-Aldrich) were captured using a confocal laser scanning microscope (FV1000-D; Olympus) or a BX51 fluorescence microscope (Olympus). The laser for detection of GFP and AF488 fluorescence was used as the excitation source at 488 nm, and fluorescence was recorded between 515 and 545 nm. The laser for detection of Calcofluor white and aniline blue was used as the excitation source at 405 nm, and fluorescence was recorded between 425 and 475 nm. Aniline blue/WGA-AF488 staining was performed according to Takemoto et al. (2011).

### 4.7 Yeast Two-Hybrid Assay

Yeast two-hybrid assays were performed according to Takemoto et al. (2011). The yeast strain AH109 was co-transformed with prey (pGADT7 derivatives) and bait (pGBKT7 derivatives) plasmids (Table S4) using the *S. cerevisiae* Direct Transformation Kit (Wako, Japan). Yeast strains used in this study are listed in Table S6. Transformants were selected on SD medium lacking leucine and tryptophan (-L/-T). For interaction assays, transformants were plated on SD medium lacking leucine and tryptophan (-L/-T) or lacking leucine, tryptophan, histidine, and adenine (-L/-T/-H/-A). Growth on the latter medium was interpreted as evidence of interaction between bait and prey proteins.

### 4.8 Bioinformatics and Structural Prediction and Visualization of the Cdc25-RasB Complex

Sequence data were analyzed and annotated using MacVector (version 18.8.2 or earlier; MacVector Inc., USA). The deduced protein sequences of fungal Cdc25 and RasB small GTPases (Figure S1 and S3) were aligned using the ClustalW program (Thompson et al., 1994) with default parameters. Phylogenetic analysis was conducted using the neighbor-joining method (Saitou and Nei, 1987). Domain annotation of Cdc25 and RasB was performed using InterProScan (Mulder and Apweiler, 2008) and the NCBI Conserved Domain Database (CDD).

The three-dimensional structure of the *E. festucae* Cdc25–RasB complex was predicted using AlphaFold3 (Abramson et al., 2024) implemented in the AlphaFold Server (https://alphafoldserver.com). Full-length amino acid sequences of Cdc25 and RasB were submitted in multimer mode with default parameters. Five structural models were generated, and confidence metrics including predicted Local Distance Difference Test (pLDDT), Predicted Aligned Error (PAE), interface predicted TM (ipTM), and predicted TM (pTM) scores were extracted from the AlphaFold output (Table S1). The structural visualization shown in Figure 5c was obtained from the AlphaFold Server output.

## Author Contributions

DT designed the research. MI, SK, AO, AM, YK, AT, and DT conducted the experiments. MI, SK, AO, YK, AT, and DT analyzed the data, DT supervised the experiments. DT wrote and edited the manuscript.

## Acknowledgements

We thank Emeritus Professor Barry Scott (Massey University, New Zealand) for providing *E. festucae* strain Fl1. We also thank Professor Kazuhito Kawakita, Dr. Sotora Chiba and Dr. Ikuo Sato (Nagoya University, Japan) for valuable suggestions, and the Radioisotope Research Center, Nagoya University, for technical assistance. This work was supported in part by the Novartis Foundation Japan for the Promotion of Science, Toyoaki Scholarship Foundation, a Grants-in-Aid for Scientific Research on Innovative Areas “Signaling functions of reactive oxygen species” (No. 23117719) from MEXT, Japan, and a Grant-in-Aid for Scientific Research (C) Generative Research Fields (No. 16KT0145) from the Japan Society for the Promotion of Science (JSPS). The authors used ChatGPT (OpenAI, GPT-5, 2025) only for English language editing of the manuscript. The authors verified and edited all AI-assisted text, and take full responsibility for the final content.

## Data Availability Statement

All data generated and analyzed during this study are included in this published article and its Supporting Information files.

## Supporting Information

**Figure S1** Alignment of the deduced amino acid sequence of EfCdc25 with Cdc25 from fungal species.

**Figure S2** Characterization of *E. festucae cdc25* mutant complemented strains.

**Figure S3** Alignment of the deduced amino acid sequence of EfRasB with RasB from fungal species.

**Figure S4** Subcellular localization of GFP-Cdc25 in hyphae of *E. festucae*.

**Figure S5** Attempted disruption of *rasB* gene in *E. festucae*.

**Figure S6** Effect of overexpressing constitutively active RasB on hyphal morphology of *E. festucae*.

**Figure S7** Strategy for deletion of *E. festucae so* gene.

**Table S1** Summary of AlphaFold3-predicted *E. festucae* Cdc25–RasB interfaces.

**Table S2** *Epichloë festucae* strains used in this study

**Table S3** Vectors for *Epichloë festucae* transformation used in this study

**Table S4** Vectors for yeast transformation used in this study

**Table S5** Primers for vector construction used in this study

**Table S6** Yeast strains used in this study

## References

1. Abramson, J., Adler, J., Dunger, J., Evans, R., Green, T., Pritzel, A., Ronneberger, O., Willmore, L., Ballard, A. J., Bambrick, J., Bodenstein, S. W., Hung, C.-C., O’Neill, M., Reiman, D., Tunyasuvunakool, K., Wu, Z., Žemgulytė, A., Arvaniti, E., Beattie, C., Bertolli, O., Bridgland, A., Cherepanov, A., Congreve, M., Cowen-Rivers, A. I., Cowie, A., Figurnov, M., Fuchs, F. B., Gladman, H., Jain, R., Khan, Y. A., Low, C. M. R., Perlin, K., Potapenko, A., Savy, P., Singh, S., Stecula, A., Thillaisundaram, A., Tong, C., Yakneen, S., Zhong, E. D., Zielinski, M., Žídek, A., Bapst, V., Kohli, P., Jaderberg, M., Hassabis, D. and Jumper, J. 2024. Accurate structure prediction of biomolecular interactions with AlphaFold 3. Nature 630: 493–500. 10.1038/s41586-024-07487-w

2. Becker, Y., Eaton, C.J., Brasell, E., May, K.J., Becker, M., Hassing, B., Cartwright, G.M., Reinhold, L., and Scott, B. 2015. The fungal cell-wall integrity MAPK cascade is crucial for hyphal network formation and maintenance of restrictive growth of *Epichloë festucae* in symbiosis with *Lolium perenne*. Molecular Plant-Microbe Interactions 28: 69–85. 10.1094/MPMI-06-14-0183-R

3. Bush, L.P., Wilkinson, H.H. and Schardl, C.L. 1997. Biotic interactions between endophytic fungi and their host grasses. Plant Physiology 114: 1–7. 10.1104/pp.114.1.1

4. Byrd, A.D., Schardl, C.L., Songlin, P.J., Mogen, K.L. and Siegel, M.R. 1990. The *β*-tubulin gene of *Epichloë typhina* from perennial ryegrass (*Lolium perenne*). Current Genetics 18: 347–354. 10.1007/BF00318216

5. Charlton, N.D., Shoji, J.-Y., Ghimire, S.R., Nakashima, J. and Craven, K.D. 2012. Deletion of the fungal gene soft disrupts the mutualistic symbiosis between the grass endophyte *Epichloë festucae* and the host plant. Eukaryotic Cell 11: 1463–1471. 10.1128/EC.00107-12

6. Chen, A., Ju, Z., Wang, J., Wang, J., Wang, H., Wu, J., Yin, Y., Zhao, Y., Ma, Z. and Chen, Y. 2020. The RasGEF FgCdc25 regulates fungal development and virulence in *Fusarium graminearum* via cAMP and MAPK signalling pathways. Environmental Microbiology 22: 5109–5124. 10.1111/1462-2920.15230

7. Christensen, M.J., Bennett, R.J., Ansari, H.A., Koga, H., Johnson, R.D., Bryan, G.T., Simpson, W.R., Koolaard, J.P. Nickless, E.M. and Voisey C.R. 2008. *Epichloë* endophytes grow by intercalary hyphal extension in elongating grass leaves. Fungal Genetics and Biology 45: 84–93. 10.1016/j.fgb.2007.07.013

8. Fischer, M.S. and Glass, N.L. 2019. Communicate and fuse: how filamentous fungi establish and maintain an interconnected mycelial network. Frontiers in Microbiology 10: 619. 10.3389/fmicb.2019.00619

9. Fortwendel, J.R. 2015. Orchestration of morphogenesis in filamentous fungi: conserved roles for Ras signaling networks. Fungal Biology Reviews 29: 54–62. 10.1016/j.fbr.2015.04.003

10. Fu, C., Iyer, P., Herkal, A., Abdullah, J., Stout, A. and Free, S.J. 2011. Identification and characterization of genes required for cell-to-cell fusion in *Neurospora crassa*. Eukaryotic Cell 10: 1100–1109. 10.1128/EC.05003-11

11. Green, K.A., Becker, Y., Tanaka, A., Takemoto, D., Fitzsimons, H.L., Seiler, S., Lalucque, H., Silar, P. and Scott, B. 2017. SymB and SymC, two membrane-associated proteins, are required for *Epichloë festucae* hyphal cell–cell fusion and maintenance of a mutualistic interaction with *Lolium perenne*. Molecular Microbiology 103: 657–677. 10.1111/mmi.13580

12. Kayano, Y., Tanaka, A., Akano, F., Scott, B. and Takemoto, D. 2013. Differential roles of NADPH oxidases and associated regulators in polarized growth, conidiation and hyphal fusion in the symbiotic fungus *Epichloë festucae*. Fungal Genetics and Biology 56: 87–97. 10.1016/j.fgb.2013.05.001

13. Kayano, Y., Tanaka, A. and Takemoto, D. 2018. Two closely related Rho GTPases, Cdc42 and RacA, of the endophytic fungus *Epichloë festucae* have contrasting roles for ROS production and symbiotic infection synchronized with the host plant. PLoS Pathogens 14: e1006840. 10.1371/journal.ppat.1006840

14. Kuldau, G.A. and Bacon, C.W. 2008. Clavicipitaceous endophytes: their ability to enhance resistance of grasses to multiple stresses. Biological Control 46: 57–71. 10.1016/j.biocontrol.2008.01.023

15. Kuroyanagi, T., Bulasag, A.S., Fukushima, K., Ashida, A., Suzuki, T., Tanaka, A., Camagna, M., Sato, I., Chiba, S., Ojika, M. and Takemoto, D. 2022. *Botrytis cinerea* identifies host plants via the recognition of antifungal capsidiol to induce expression of a specific detoxification gene. PNAS Nexus 1: pgac274. 10.1093/pnasnexus/pgac274

16. Latch, G.C.M. and Christensen, M.J. 1985. Artificial infection of grasses with endophytes. Annals of Applied Biology 107: 17–24. 10.1111/j.1744-7348.1985.tb01543.x

17. Li, H., Shen, X., Wu, W., Zhang, W. and Wang, Y. 2023. Ras2 is responsible for the environmental responses, melanin metabolism, and virulence of *Botrytis cinerea*. J. Fungi (Basel) 9: 432. 10.3390/jof9040432

18. Mintzer, K.A. and Field, J. 1999. The SH3 domain of the *Saccharomyces cerevisiae* Cdc25p binds adenylyl cyclase and facilitates Ras regulation of cAMP signalling. Cellular Signalling 11: 127–135. 10.1016/S0898-6568(98)00044-8

19. Mulder, N.J. and Apweiler, R. 2008. The InterPro database and tools for protein domain analysis. Current Protocols in Bioinformatics Chapter 2: Unit 2.7. 10.1002/0471250953.bi0207s21

20. Müller, P., Katzenberger, J.D., Loubradou, G. and Kahmann, R. 2003. Guanyl nucleotide exchange factor Sql2 and Ras2 regulate filamentous growth in *Ustilago maydis*. Eukaryotic Cell 2: 609–617. 10.1128/EC.2.3.609-617

21. Niones, J.T. and Takemoto, D. 2015. VibA, a homologue of a transcription factor for fungal heterokaryon incompatibility, is involved in antifungal compound production in the plant-symbiotic fungus *Epichloë festucae*. Eukaryotic Cell 14: 13–24. 10.1128/EC.00034-14

22. Nowrousian, M., Frank, S., Koers, S., Strauch, P., Weitner, T., Ringelberg, C., Dunlap, J.C., Loros, J.J., and Kück, U. 2007. The novel ER membrane protein PRO41 is essential for sexual development in the filamentous fungus *Sordaria macrospora*. Molecular Microbiology 64(4): 923–937. 10.1111/j.1365-2958.2007.05694.x

23. Park, G., Xue, C., Zhao, X., Kim, Y., Orbach, M. and Xu, J.R. 2006. Multiple upstream signals converge on the adaptor protein Mst50 in *Magnaporthe grisea*. Plant Cell 18: 2822–2835. 10.1105/tpc.105.038422

24. Rowan, D.D., Hunt, M.B. and Gaynor, D.L. 1986. Peramine, a novel insect feeding deterrent from ryegrass infected with the endophyte *Acremonium loliae*. Journal of the Chemical Society, Chemical Communications 1: 935–936. 10.1039/C39860000935

25. Saikia, S., Takemoto, D., Tapper, B.A., Lane, G.A., Fraser, K. and Scott, B. 2012. Functional analysis of an indole-diterpene gene cluster for lolitrem B biosynthesis in the grass endosymbiont *Epichloë festucae*. FEBS Letters 586: 2563–2569. 10.1016/j.febslet.2012.06.035

26. Saitou, N. and Nei, M. 1987. The neighbor-joining method: a new method for reconstructing phylogenetic trees. Molecular Biology and Evolution 4: 406–425. 10.1093/oxfordjournals.molbev.a040454

27. Sánchez, O., Navarro, R.E. and Aguirre, J. 1998. Increased transformation frequency and tagging of developmental genes in *Aspergillus nidulans* by restriction enzyme-mediated integration (REMI). Molecular and General Genetics 258: 89–94. 10.1007/s004380050710

28. Schardl, C.L., Leuchtmann, A. and Spiering, M.J. 2004. Symbioses of grasses with seedborne fungal endophytes. Annual Review of Plant Biology 55: 315–340. 10.1146/annurev.arplant.55.031903.141735

29. Takemoto, D., Tanaka, A. and Scott, B. 2006a. A p67^Phox^-like regulator is recruited to control hyphal branching in a fungal–grass mutualistic symbiosis. Plant Cell 18: 2807–2821. 10.1105/tpc.106.046169

30. Takemoto, D., Tanaka, A. and Scott, B. 2006b. NADPH oxidases in fungi: diverse roles of reactive oxygen species in fungal cellular differentiation. Fungal Genetics and Biology 43: 1017–1026. 10.1016/j.fgb.2006.06.008

31. Takemoto, D., Kamakura, S., Saikia, S., Becker, Y., Wrenn, R., Tanaka, A., Sumimoto, H. and Scott, B. 2011. Polarity proteins Bem1 and Cdc24 are components of the filamentous fungal NADPH oxidase complex. Proceedings of the National Academy of Sciences of the United States of America 108: 2861–2866. 10.1073/pnas.1017309108

32. Tanaka, A., Tapper, B.A., Popay, A., Parker, E.J. and Scott, B. 2005. A symbiosis-expressed nonribosomal peptide synthetase from a mutualistic fungal endophyte of perennial ryegrass confers protection to the symbiotum from insect herbivory. Molecular Microbiology 57: 1036–1050. 10.1111/j.1365-2958.2005.04747.x

33. Tanaka, A., Christensen, M.J., Takemoto, D., Park, P. and Scott, B. 2006. Reactive oxygen species play a role in regulating a fungus-perennial rygrass mutualistic interaction. Plant Cell 18: 1052–1066. 10.1105/tpc.105.039263

34. Tanaka, A., Takemoto, D., Hyon, G.S., Park, P. and Scott, B. 2008. NoxA activation by the small GTPase RacA is required to maintain a mutualistic symbiotic association between *Epichloë festucae* and perennial ryegrass. Molecular Microbiology 68: 1165–1178. 10.1111/j.1365-2958.2008.06217.x

35. Tanaka, A., Takemoto, D., Chujo, T. and Scott, B. 2012. Fungal endophytes of grasses. Current Opinion in Plant Biology 15: 462–468. 10.1016/j.pbi.2012.03.007

36. Tanaka, A., Cartwright, G.M., Saikia, S., Kayano, Y., Takemoto, D., Kato, M., Tsuge, T. and Scott, B. 2013. ProA, a transcriptional regulator of fungal fruiting body development, regulates leaf hyphal network development in the *Epichloë festucae*–*Lolium perenne* symbiosis. Molecular Microbiology 90: 551–568. 10.1111/mmi.12385

37. Tanaka, A., Kamiya, S., Ozaki, Y., Kameoka, S., Kayano, Y., Saikia, S., Akano, F., Uemura, A., Takagi, H., Terauchi, R., Maruyama, J.I., Hammadeh, H.H., Fleissner, A., Scott, B. and Takemoto, D. 2020. A nuclear protein NsiA from *Epichloë festucae* interacts with a MAP kinase MpkB and regulates the expression of genes required for symbiotic infection and hyphal cell fusion. Molecular Microbiology 114: 626–640. 10.1111/mmi.14568

38. Thompson, J.D., Higgins, D.G. and Gibson, T.J. 1994. CLUSTAL W: improving the sensitivity of progressive multiple sequence alignment through sequence weighting, position-specific gap penalties and weight matrix choice. Nucleic Acids Research 22: 4673–4680. 10.1093/nar/22.22.4673

39. Turrà, D., Segorbe, D. and Di Pietro, A. 2014. Protein kinases in plant-pathogenic fungi: conserved regulators of infection. Annual Review of Phytopathology 52: 267–288. 10.1146/annurev-phyto-102313-050143

40. van der Lee, R., Buljan, M., Lang, B., Weatheritt, R.J., Daughdrill, G.W., Dunker, A.K., Fuxreiter, M., Gough, J., Gsponer, J., Jones, D.T., Kim, P.M., Kriwacki, R.W., Oldfield, C.J., Pappu, R.V., Tompa, P., Uversky, V.N., Wright, P.E. and Babu, M.M. 2014. Classification of intrinsically disordered regions and proteins. Chemical Reviews 114: 6589–6631. 10.1021/cr400525m

41. Voisey, C.R., Christensen, M.T., Johnson, L.J., Forester, N.T., Gagic, M., Bryan, G.T., Simpson, W.R., Fleetwood, D.J., Card, S.D., Koolaard, J.P., Maclean, P.H. and Johnson, R.D. 2016. cAMP signaling regulates synchronised growth of symbiotic *Epichloë* fungi with the host grass *Lolium perenne*. Frontiers in Plant Science 7: 1546. 10.3389/fpls.2016.01546

42. Wilkinson, H.H., Siegel, M.R., Blankenship, J.D., Mallory, A.C., Bush, L.P. and Schardl, C.L. 2000. Contribution of fungal loline alkaloids to protection from aphids in a grass–endophyte mutualism. Molecular Plant-Microbe Interactions 13: 1027–1033. 10.1094/MPMI.2000.13.10.1027

43. Yan, Y., Tang, J., Yuan, Q., Gu, Q., Liu, H., Huang, J., Hsiang, T. and Zheng, L. 2020. ChCDC25 regulates infection-related morphogenesis and pathogenicity of the crucifer anthracnose fungus *Colletotrichum higginsianum*. Frontiers in Microbiology 11: 763. 10.3389/fmicb.2020.00763

