## Supplemental Figures S1-S7 for "Cdc25-Mediated Activation of the Small GTPase RasB Is Essential for Hyphal Fusion and Symbiotic Infection of *Epichloë festucae*"

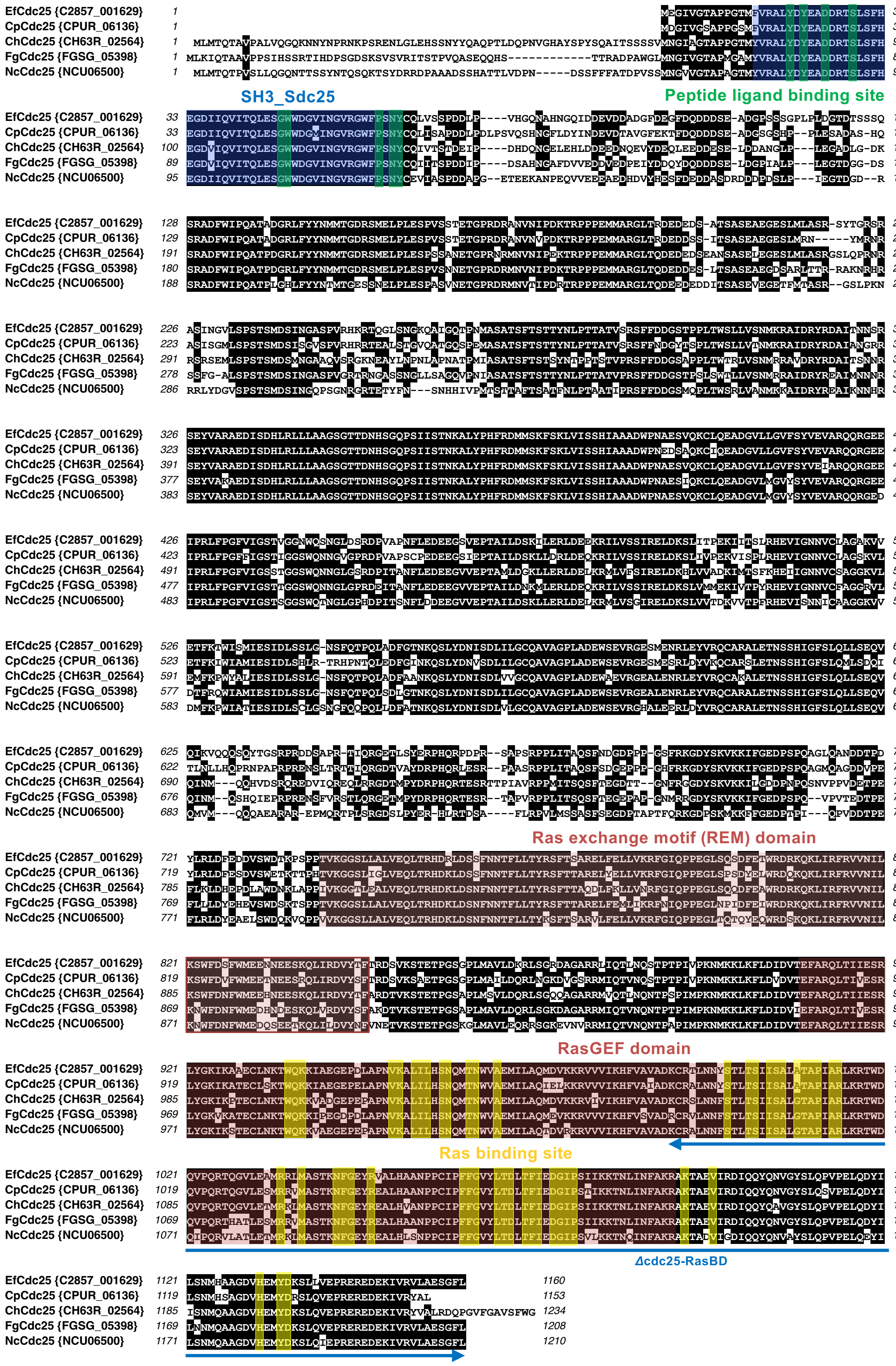


**FIGURE S1** | Alignment of the deduced amino acid sequence of *Epichloë festucae* Cdc25 (EfCdc25; C2857_001629) with Cdc25 from *Claviceps purpurea* (CpCdc25; CPUR_06136), *Colletotrichum higginsianum* (ChCdc25; CH63R_02564), *Fusarium graminearum* (FgCdc25; FGSG_05398) and *Neurospora crassa* (NcCdc25; NUC06500). Src homology 3 (SH3) domain with a peptide ligand-binding site (SH3_Sdc25, blue), Ras exchange motif (REM) and Ras guanine nucleotide exchange factor (RasGEF) domains (pink) with conserved Ras-binding sites (yellow) are indicated. The region deleted in the *Δcdc25-RasBD* strains corresponds to the C-terminal portion of the RasGEF and Ras-binding domain is indicated by the horizontal blue double-headed arrow.

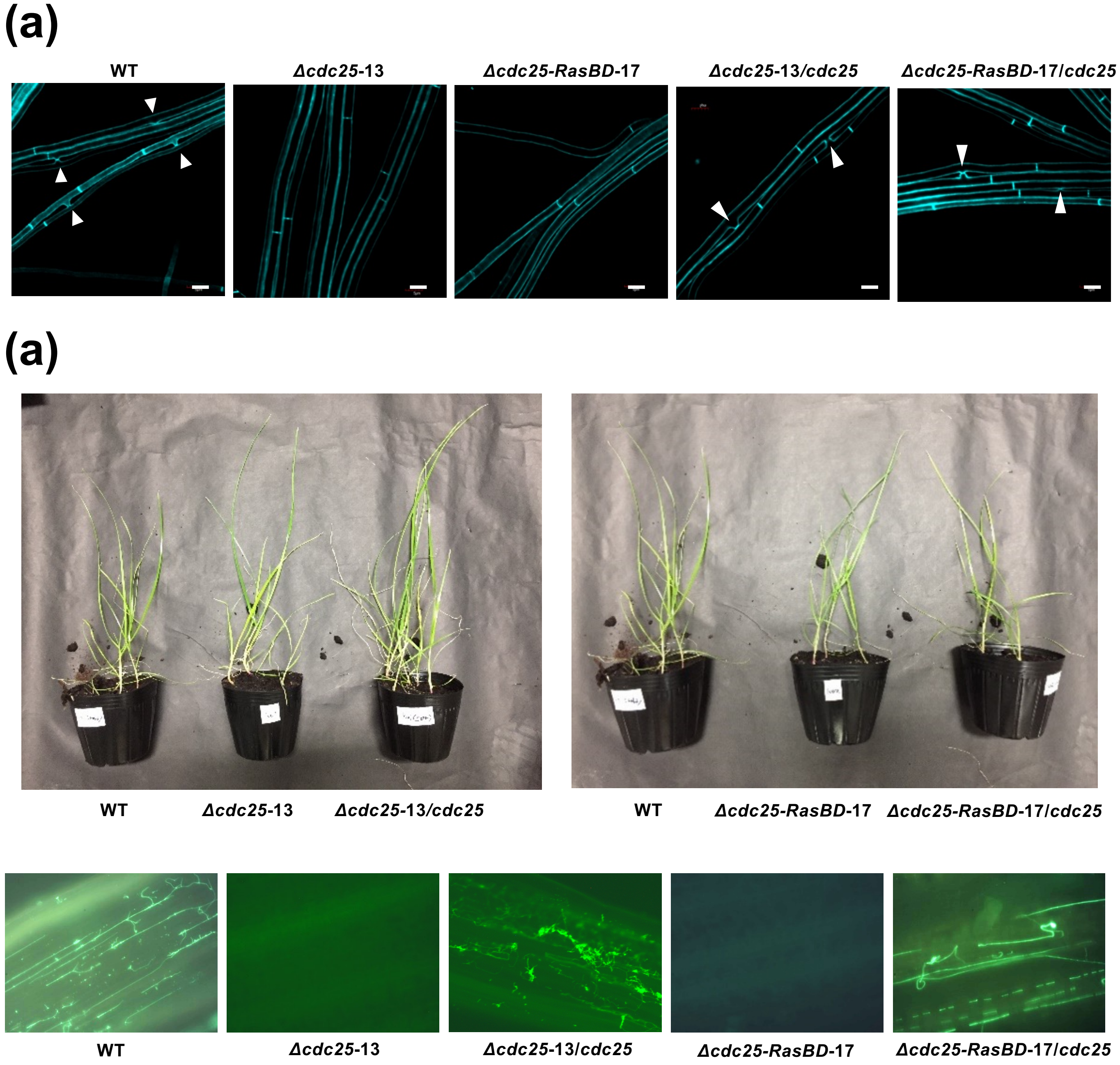


**FIGURE S2 | (a)** Hyphal fusion of *Epichloë festucae* wild type (WT), *cdc25* mutants, and complemented strains. Hyphal morphology and fusion were examined in *E. festucae* strains grown on water agar, stained with Calcofluor white, and observed using confocal laser scanning microscopy. Arrowheads indicate hyphal fusion events. Bars = 5 µm. **(b)** Infection of *E. festucae cdc25* mutants and complemented strains in perennial ryegrass. (top) Perennial ryegrass plants infected with *E. festucae* WT, *cdc25* mutants, and complemented strains. (bottom) Hyphae of *E. festucae* WT, *cdc25* mutants, and complemented strains within host tissues. Leaves inoculated with each strain were stained with aniline blue and observed under fluorescence microscopy.

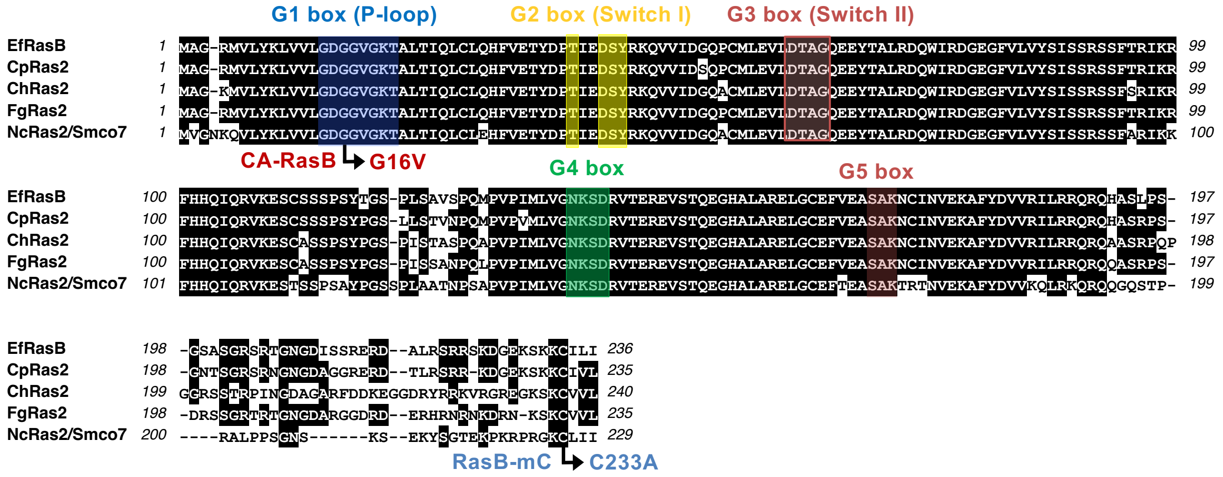


**FIGURE S3 |** Alignment of the deduced amino acid sequence of *Epichloë festucae* RasB (EfRasB) with RasB/Ras2 from *Claviceps purpurea* (CpRas2), *Colletotrichum higginsianum* (ChRas2), *Fusarium graminearum* (FgRas2), and *Neurospora crassa* (NcRas2/Smco7). Conserved domains among Ras GTPases are boxed. Amino acid substitutions introduced to generate the constitutively active (CA) form and the mutation at the geranylgeranylation site used for the yeast two-hybrid assay are indicated.

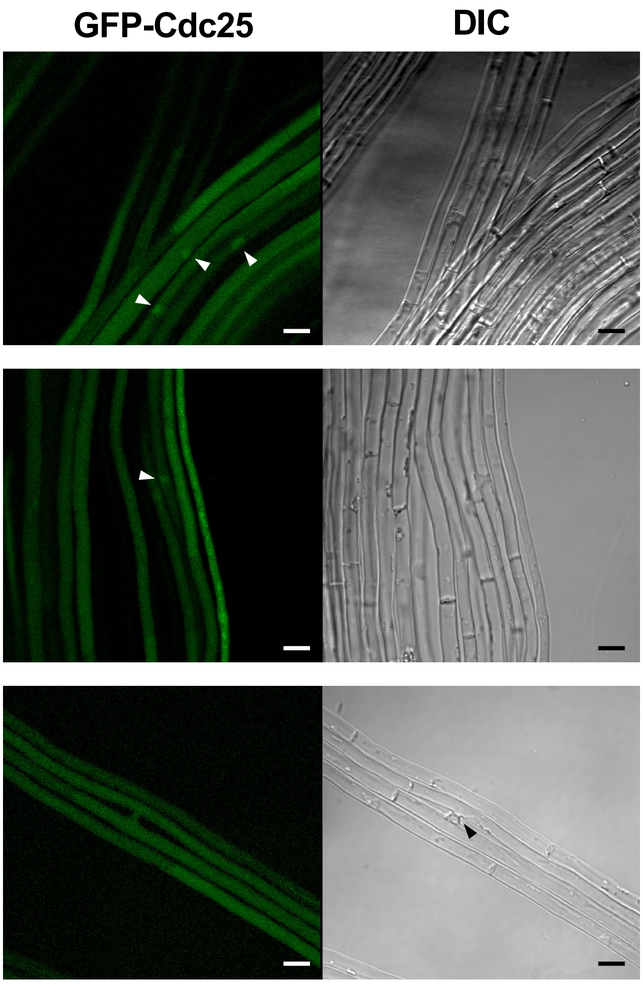


**FIGURE S4** | Subcellular localization of GFP-Cdc25 in hyphae of *Epichloë festucae*. GFP-RasB was expressed under the control of the *TEF* promoter. Localization of GFP-Cdc25 was examined in three individual transformants. White and black arrowheads indicate the potential local accumulation of GFP-Cdc25 and the sites of hyphal fusion, respectively. Bars = 5 μm.

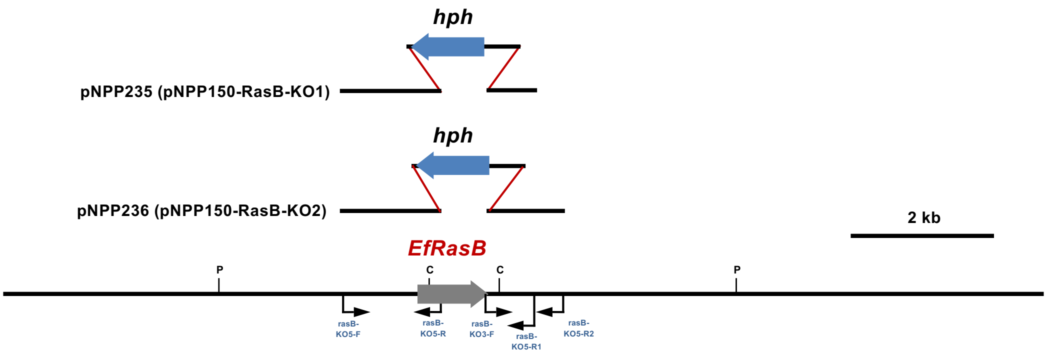


**FIGURE S5** | Attempted disruption of *rasB* gene in *Epichloë festucae*.
Physical map of the *rasB* wild-type genomic region and linear inserts of *rasB* replacement constructs pNPP150-RasB-KO1 and pNPP150-RasB-KO2. P, *Pst*I.

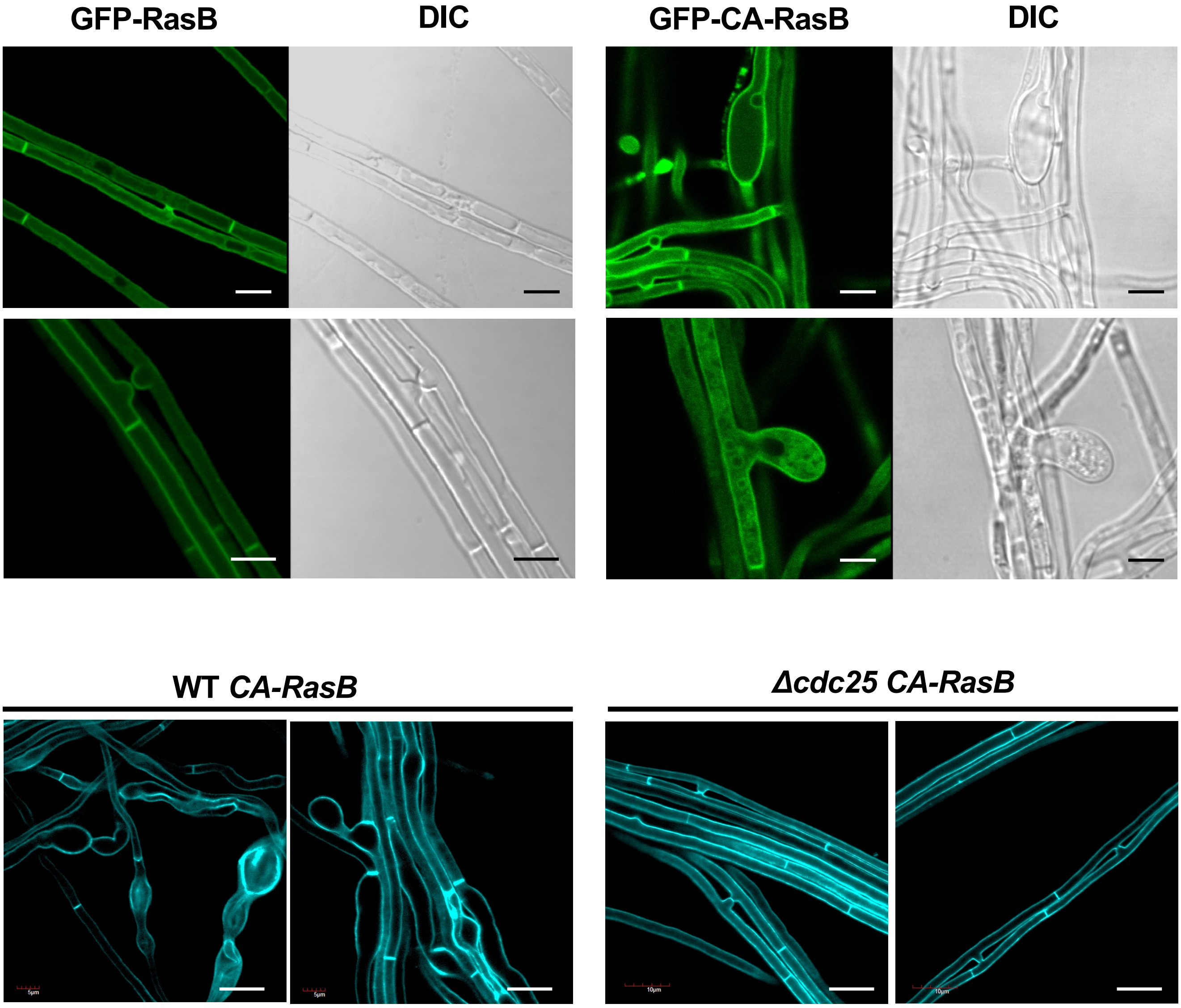


**FIGURE S6** | Effect of overexpressing constitutively active RasB on hyphal morphology of *Epichloë festucae*.

**(a)** GFP-RasB or GFP-CA-RasB was expressed under the control of the *TEF* promoter. Localization of GFP fluorescence and hyphal morphology were observed in hyphae grown on water agar. Bars = 5 µm. **(b)** Hyphal morphology of *E. festucae* wild type (WT) or *Δcdc25* expressing CA-RasB under the control of *TEF* promoter. *E. festucae* transformants were grown on water agar, stained with Calcofluor white, and observed using confocal laser scanning microscopy. Bars = 10 µm.

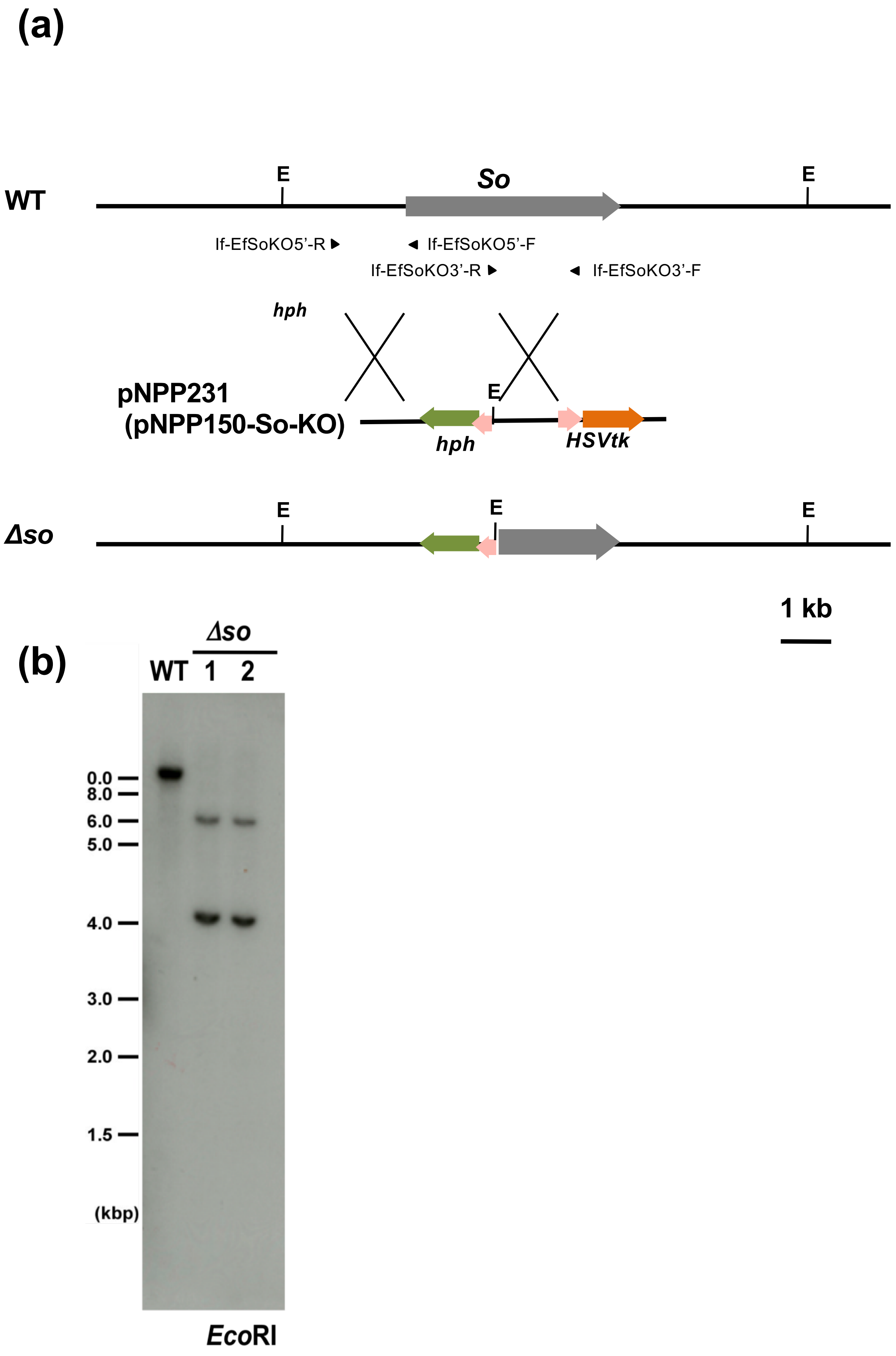


**FIGURE S7** | Strategy for deletion of *Epichloë festucae so* gene.
**(a)** Physical map of the *so* wild-type genomic region and linear insert of *so* replacement construct pNPP231 (pNPP150-so-KO). E, *Eco*RI. **(b)** Autoradiograph of Southern blot of *Eco*RI-digested genomic DNA of *E. festucae* wild-type (WT) and *Δso* with [$^{32}$P]-labeled pNPP231. The sizes of marker DNA fragments are given in kb.
