## Supplemental Tables S1-S6 for "Cdc25-Mediated Activation of the Small GTPase RasB Is Essential for Hyphal Fusion and Symbiotic Infection of *Epichloë festucae*"

**Table S1.** Summary of AlphaFold3-predicted *Epichloë festucae* Cdc25–RasB interfaces and domain architecture

| Model | Mean pLDDT | ipTM | pTM | Rank score | Mean PAE (Å) | Interface_residues (EfCdc25/RasB) | Interface_type | Major domains in EfCdc25 | Notes | Recycles |
|---|---|---|---|---|---|---|---|---|---|---|
| 0 | 85.1 | 0.77 | 0.51 | 0.84 | 7.6 | 890-1070 / 35-65 | RasGEF-Switch I/II | SH3, REM, RasGEF | Top-ranked model; stable interface | 10 |
| 1 | 85.0 | 0.76 | 0.51 | 0.83 | 7.8 | 895-1065 / 38-67 | RasGEF-Switch I/II | SH3, REM, RasGEF | Similar to model 0, minor RMSD shift | 10 |
| 2 | 85.2 | 0.76 | 0.51 | 0.83 | 7.5 | 880-1080 / 33-70 | RasGEF-Switch I/II | SH3, REM, RasGEF | Consistent binding orientation | 10 |
| 3 | 84.7 | 0.76 | 0.51 | 0.83 | 7.9 | 900-1075 / 30-68 | RasGEF-Switch I/II | SH3, REM, RasGEF | Slightly looser interface | 10 |
| 4 | 84.9 | 0.76 | 0.51 | 0.82 | 8 | 895-1070 / 34-66 | RasGEF-Switch I/II | SH3, REM, RasGEF | Lower rank; similar topology | 10 |

pLDDT, predicted Local Distance Difference Test; ipTM, interface predicted Template Modelling; pTM, predicted Template Modelling; PAE, Predicted Aligned Error; RMSD, Root Mean Square Deviation.

**Table S2.** *Epichloë festucae* strains used in this study.

| Fungal strains | Relevant characteristics | Reference |
|---|---|---|
| ***Epichloë festucae*** | | |
| Fl1 | Wild type | Young *et al.* 2005 |
| RPA41 | Fl1/pNPP1; $Hyg^R$ | This study |
| RPA112 | Fl1/pNPP1; $Hyg^R$ | This study |
| RPA519 | Fl1/pNPP1; $Hyg^R$ | This study |
| *Δham8*-1 | Fl1/*Δham8*::P*trpC*-*hph*; $Hyg^R$ | Tanaka *et al.* 2020 |
| *ΔmpkB*-1 | Fl1/*ΔmpkB*::P*trpC*-*hph*; $Hyg^R$ | Tanaka et al. 2020 |
| *Δso* | Fl1/*Δso*::P*trpC*-*hph* (pNPP231); $Hyg^R$ | This study |
| *ΔnoxA* | Fl1/*ΔnoxA*::P*trpC*-*hph*; $Hyg^R$ | Tanaka *et al.* 2006 |
| *Δcdc25-13* | Fl1/*Δcdc25*::P*trpC*-*hph* (pNPP233); $Hyg^R$ | This study |
| *Δcdc25-RasBD-16* | Fl1/*Δcdc25-RasBD*::P*trpC*-*hph* (pNPP234); $Hyg^R$ | This study |
| *Δcdc25-RasBD-17* | Fl1/*Δcdc25-RasBD*::P*trpC*-*hph* (pNPP234); $Hyg^R$ | This study |
| *Δcdc25-RasBD-18* | Fl1/*Δcdc25-RasBD*::P*trpC*-*hph* (pNPP234); $Hyg^R$ | This study |
| *Δcdc25-RasBD-25* | Fl1/*Δcdc25-RasBD*::PtrpC-hph (pNPP234); $Hyg^R$ | This study |
| *Δcdc25-RasBD-26* | Fl1/*Δcdc25-RasBD*::P*trpC*-*hph* (pNPP234); $Hyg^R$ | This study |
| *Δcdc25-13-GFP* | *Δcdc25-13*/pNPP99; $Hyg^R$, $Gen^R$ | This study |
| *Δcdc25-RasBD-17-GFP* | *Δcdc25-RasBD-17*/pNPP99; $Hyg^R$, $Gen^R$ | This study |
| *Δcdc25-13/cdc25* | *Δcdc25-13*/pNPP241; $Hyg^R$, $Gen^R$ | This study |
| *Δcdc25-RasBD-17/cdc25* | *Δcdc25-RasBD-17*/pNPP241; $Hyg^R$, $Gen^R$ | This study |
| *GFP-RasB* | Fl1/pNPP223; $Hyg^R$ | This study |
| *GFP-CA-RasB* | Fl1/pNPP224; $Hyg^R$ | This study |
| *GFP-Cdc25* | Fl1/pNPP222; $Hyg^R$ | This study |
| *CA-RasB* | Fl1/pNPP225; $Hyg^R$ | This study |
| *Δcdc25-13/CA-RasB* | *Δcdc25-13*/pNPP225, pSF17; $Hyg^R$, $Gen^R$ | This study |
| *Δcdc25-RasBD-17/CA-RasB* | *Δcdc25-RasBD-17*/pNPP225, pSF17; $Hyg^R$, $Gen^R$ | This study |
| *ΔmpkB/CA-RasB* | *ΔmpkB*/pNPP225, pSF17; $Hyg^R$, $Gen^R$ | This study |
| *ΔnoxA/CA-RasB* | *ΔnoxA*/pNPP225, pSF17; $Hyg^R$, $Gen^R$ | This study |

**Table S3.** Plasmids for *Epichloë festucae* transformation used in this study

| Vector name | Base vector | Insert | Primers used to amplify insert | References | Description |
|---|---|---|---|---|---|
| **Base vectors** | | | | | |
| pNPP1 | - | - | - | Kayano et al. 2013 | Vector used for REMI, $Amp^R$/$Hyg^R$, |
| pPN94 | - | - | - | Takemoto et al. 2006 | Base vector for gene expression under TEF promoter, $Amp^R$/$Hyg^R$ |
| pSF17.1 | - | - | - | Tanaka et al. 2008 | Base vector for complementation and co-transformation, $Amp^R$/$Gen^R$ |
| pNPP141 (pPN94-GFP-3GA) | - | - | - | Kayano et al. 2013 | Vector for expression of GFP-tagged protein, $Amp^R$/$Hyg^R$ |
| pNPP150 | - | - | - | Niones and Takemoto 2015 | Base vector for KO (HSVtk), $Amp^R$/$Hyg^R$ |
| pNPP99 | - | - | - | Kayano et al. 2013 | Expression of GFP, $Amp^R$/$Gen^R$ |
| **Plasmids for gene knock out and complementation** | | | | | |
| pNPP231 (pNPP150-So-KO) | pNPP150 | 5'So-PtrpC-hph-3'So | IF So-KO5-F, IF So-KO5-R, IF So-KO3-F, IF So-KO3-R | This study | So KO, $Amp^R$/$Hyg^R$ |
| pNPP232 (pNPP150-RPA112-KO) | pNPP150 | 5'C2857_000796-PtrpC-hph-3'C2857_000796 | IF RPA112-KO5-F, IF RPA112-KO5-R, IF RPA112-KO3-F, IF RPA112-KO3-R | This study | C2857_000796 KO, $Amp^R$/$Hyg^R$ |
| pNPP233 (pNPP150-Cdc25-KO) | pNPP150 | 5'Cdc25-PtrpC-hph-3'Cdc25 | IF Cdc25-KO5-F1, IF Cdc25-KO5-R1, IF Cdc25-KO3-F, IF Cdc25-KO3-R | This study | Cdc25 KO, $Amp^R$/$Hyg^R$ |
| pNPP234 (pNPP150-Cdc25-RasBD-KO) | pNPP150 | 5'Cdc25RasBD-PtrpC-hph-3'Cdc25 | IF Cdc25-KO5-F2, IF Cdc25-KO5-R2, IF Cdc25-KO3-F, IF Cdc25-KO3-R | This study | Cdc25 Ras binding domain KO, $Amp^R$/$Hyg^R$ |
| pNPP235 (pNPP150-RasB-KO1) | pNPP150 | 5'RasB-PtrpC-hph-3'RasB | IF RasB-KO5-F, IF RasB-KO5-R, IF RasB-KO3-F1, IF RasB-KO3-R1 | This study | RasB KO, $Amp^R$/$Hyg^R$ |
| pNPP236 (pNPP150-RasB-KO2) | pNPP150 | 5'RasB-PtrpC-hph-3'RasB2 | IF RasB-KO5-F, IF RasB-KO5-R, IF RasB-KO3-F2, IF RasB-KO3-R2 | This study | RasB KO, $Amp^R$/$Hyg^R$ |
| pNPP241 (pSF17-Cdc25) | pSF17.1 | *Cdc25* | pSF17-cdc25-F2, pSF17-cdc25-R2 | This study | Complementation of *cdc25* , $Amp^R$/$Gen^R$ |
| **Plasmids for gene expression in *E. festucae*** | | | | | |
| pNPP222 (pPN94-GFP-Cdc25) | pNPP141 | GFP-Cdc25 | IF pPN94-GFP-3GA-Cdc25-F, IF pPN94-GFP-3GA-Cdc25-R | This study | Expression of GFP-Cdc25, $Amp^R$/$Hyg^R$ |
| pNPP223 (pPN94-GFP-RasB) | pNPP141 | GFP-RasB | IF pPN94-GFP-3GA-RasB-F, IF pPN94-GFP-3GA-RasB-R | This study | Expression of GFP-RasB, $Amp^R$/$Hyg^R$ |
| pNPP224 (pPN94-GFP-CA-RasB) | pNPP141 | GFP-CA-RasB | IF pPN94-GFP-3GA-RasB-F, IF pPN94-GFP-3GA-RasB-R | This study | Expression of GFP-CA-RasB, $Amp^R$/$Hyg^R$ |
| pNPP225 (pPN94-CA-RasB) | pPN94 | CA-RasB | IF pPN94-RasB-F, IF pPN94-RasB-R, IF RasB-DA-F, IF RasB-DA-F2, IF RasB-DA-F3, IF RasB-DA-R | This study | Expression of CA-RasB, $Amp^R$/$Hyg^R$ |

**Table S4.** Plasmids for yeast two-hybrid assay used in this study.

| Plasmid name | Base vector | Insert | Primers used to amplify insert | References | Description |
|---|---|---|---|---|---|
| **Base vectors** | | | | | |
| pGADT7 | - | - | - | Clontech | $Amp^R$/LEU2 |
| pGBKT7 | - | - | - | Clontech | $Kan^R$/TRP1 |
| **Plasmids for yeast two-hybrid assay** | | | | | |
| pNPP216 (pGADT7 Cdc25) | pGADT7 | Cdc25 | IF-pGADT7-Cdc25-F, IF-pGADT7-Cdc25-R | This study | Full length Cdc25 |
| pNPP217 (pGBKT7 RasA-mC) | pGBKT7 | RasA-mC | IF-pGBKT7-RasA-F, IF-pGBKT7-RasAmC-R | This study | Full length RasA with mutation in CaaX motif (C213A) |
| pNPP218 (pGBKT7 RasB-mC) | pGBKT7 | RasB-mC | IF-pGBKT7-RasB-F, IF-pGBKT7-RasBmC-R | This study | Full length RasB with mutation in CaaX motif (C233A) |
| pNPP219 (pGBKT7 RasC-mC) | pGBKT7 | RasC-mC | IF-pGBKT7-RasC-F, IF-pGBKT7-RasCmC-R | This study | Full length RasC with mutation in CaaX motif (C267A) |
| pNPP220 (pGBKT7 RhbA-mC) | pGBKT7 | RhbA-mC | IF-pGBKT7-RhbA-F, IF-pGBKT7-RhbAmC-R | This study | Full length RhbA with mutation in CaaX motif (C183A) |
| pNPP221 (pGBKT7 KrevA-mC) | pGBKT7 | KrevA-mC | IF-pGBKT7-KrevA-F, IF-pGBKT7-KrevAmC-R | This study | Full length KrevA with mutation in CaaX motif (C216A) |

**Table S5.** Primers used in this study

| Primer name | Sequence |
|---|---|
| cdc25-F | TGCGAAACGAGCAAAGACAG |
| cdc25-R | TATGCCGCATTGCTACGTAC |
| Ppro41-F | CTGAGTTGGACAAATATGGC |
| Ppro41-R | GCCACTTCGGCCTATTCTCC |
| IF So-KO5-F | TTAGGTGACACTATACATGTCAAGTCAGCG |
| IF So-KO5-R | CAGCTGCTCGAGTTCCCCCCTCTGTTTCTC |
| IF So-KO3-F | TGAGTCGTATTAATTATGGAAACCAGCGTC |
| IF So-KO3-R | AACCTGGCTTATCGATGAGAATGCTTCCCA |
| IF Cdc25-KO5-F1 | ATGCCTGCAGGTCGAATAAGTTGGTGAGACTTCCC |
| IF Cdc25-KO5-R1 | ATCCTCTAGAGTCGAGTTTGTGCCGAAGCGCACAA |
| IF Cdc25-KO5-F2 | ATGCCTGCA+C8GGTCGAGCAACTGCTCTCCGAGCAAG |
| IF Cdc25-KO5-R2 | ATCCTCTAGAGTCGAGTAGTTGTTGAGAGTTCGGC |
| IF Cdc25-KO3-F | TACCGAGCTCGAATTATCGAACCATCGGGGTCAAC |
| IF Cdc25-KO3-R | TATCATCGATGAATTTTAGATTCCATGTCGTCAGG |
| IF RPA112-KO5-F | ATGCCTGCAGGTCGACTGTGATATCAACGTCCCTG |
| IF RPA112-KO5-R | ATCCTCTAGAGTCGATTCATTTCTCGAAGCCGTCG |
| IF RPA112-KO3-F | TACCGAGCTCGAATTGAAAGACAGTCCTCGTCTTG |
| IF RPA112-KO3-R | TATCATCGATGAATTATCGGCCTTGACCAGTTCCT |
| IF-pGADT7-Cdc25-F | GGAGGCCAGTGAATTCATGGAGGGCATTGTGGGAA |
| IF-pGADT7-Cdc25-R | TCATCTGCAGCTCGAGTCATAGAAACCCGGACTCG |
| IF-pGBKT7-RasA-F | CATGGAGGCCGAATTCATGGCCGCATCCACAAAGT |
| IF-pGBKT7-RasAmC-R | GCAGGTCGACGGATCCTCACATAATGATAGCCTTG |
| IF-pGBKT7-RasB-F | CATGGAGGCCGAATTCATGGCGGGCCGTATGGTGT |
| IF-pGBKT7-RasBmC-R | GCAGGTCGACGGATCCTCATATAAGGATGGCTTTT |
| IF-pGBKT7-RasC-F | CATGGAGGCCGAATTCATGTCTTCTCAGCTGGAGC |
| IF-pGBKT7-RasCmC-R | GCAGGTCGACGGATCCTTACCAGGCCCGCAGCTTC |
| IF-pGBKT7-RhbA-F | CATGGAGGCCGAATTCATGCCTGCGCCAAAGCAGA |
| IF-pGBKT7-RhbAmC-R | GCAGGTCGACGGATCCCTACATGAGAGAGGCGTTG |
| IF-pGBKT7-KrevA-F | CATGGAGGCCGAATTCATGGCGCCTCGATTCCACG |
| IF-pGBKT7-KrevAmC-R | GCAGGTCGACGGATCCCTACAAGATTACGGCTCTG |
| IF pSF17 cdc25-F2 | GAATTCATCGATGATGCAGTTAAACTCGTTCTCGT |
| IF pSF17 cdc25-R2 | ACCGGCAGATCTGATGGGGCTGTTTTGATGAAAAG |
| IF RasB-KO5-F | ATGCCTGCAGGTCGACCAAGGGTGAGGTTGTGTAT |
| IF RasB-KO5-R | ATCCTCTAGAGTCGATGGAACCTTGCGTGCCGAAA |
| IF RasB-KO3-F1 | TACCGAGCTCGAATTCGGCAACAACGAAGCGAGTG |
| IF RasB-KO3-R1 | TATCATCGATGAATTAGGGCTGGCTGTCTGTAAGT |
| IF RasB-KO3-F2 | TACCGAGCTCGAATTACGGGGTCACCTCTTTCCGC |
| IF RasB-KO3-R2 | TATCATCGATGAATTTCATCTCTTCATGTCGACCT |
| IF pPN94-RasB-F | AACCTCTAGAGGATCATGGCGGGCCGTATGGTGTT |
| IF pPN94-RasB-R | ACGTTAAGTGCGGCCTCATATAAGGATGCATTTTT |
| IF RasB-DA-F | GACGTCGGCGTAGGAAAGAC |
| IF RasB-DA-R | TCCTACGCCGACGTCTCCCA |
| IF RasB-DA-F2 | ACAAGCTTGTGGTTCTGGGAGACGTCGGCGTAGGA |
| IF RasB-DA-F3 | GGCGGGCCGTATGGTGTTGTACAAGCTTGTGGTTC |
| IF pPN94-GFP-3GA-RasB-F | TGCTGGTGCTGAATTCATGGCGGGCCGTATGGTGT |
| IF pPN94-GFP-3GA-RasB-R | ACGTTAAGTGCGGCCTCATATAAGGATGCATTTTT |
| IF pPN94-GFP-3GA-Cdc25-F | TGCTGGTGCTGAATTCATGGAGGGCATTGTGGGAA |
| IF pPN94-GFP-3GA-Cdc25-R | ACGTTAAGTGCGGCCTCATAGAAACCCGGACTCGG |

Extension sequences for In-fusion reaction are in red letters.

Mismatches to introduce amino acid substitution are highlighted in blue letters.

Extension sequence for in-fusion reaction are shown in red letters.

**Table S6.** Yeast strains used in this study

| Yeast strains | Relevent characteristics | References |
|---|---|---|
| ***Saccharomyces cerevisiae*** | | |
| AH109 | MATa, trp1-901, Leu2-3, 112, ura3-52, his3-200, gal4⊿, gal80⊿, LYS2::GAL1UAS-GAL1TATA-HIS3, MEL1, GAL2UAS-GAL2TATA-ADE2, URA::MELIUAS-MEL1TATA-laeZ | Clontech |
| AH109 (Cdc25/empty) | AH109/pNPP216; pGBKT7; *LEU/TPR1* | This study |
| AH109 (empty/RasA) | AH109/pGADT7; pNPP217; LEU/TPR1 | This study |
| AH109 (empty/RasB) | AH109/pGADT7; pNPP218; LEU/TPR1 | This study |
| AH109 (empty/RasC) | AH109/pGADT7; pNPP219; LEU/TPR1 | This study |
| AH109 (empty/RhbA) | AH109/pGADT7; pNPP220; LEU/TPR1 | This study |
| AH109 (empty/KrevA) | AH109/pGADT7; pNPP221; LEU/TPR1 | This study |
| AH109 (Cdc25/RasA) | AH109/pNPP216; pNPP217; *LEU/TPR1* | This study |
| AH109 (Cdc25/RasB) | AH109/pNPP216; pNPP218; *LEU/TPR1* | This study |
| AH109 (Cdc25/RasC) | AH109/pNPP216; pNPP219; *LEU/TPR1* | This study |
| AH109 (Cdc25/RhbA) | AH109/pNPP216; pNPP220; *LEU/TPR1* | This study |
| AH109 (Cdc25/KrevA) | AH109/pNPP216; pNPP221; *LEU/TPR1* | This study |
